# Function and constraint in enhancers with multiple evolutionary origins

**DOI:** 10.1101/2022.01.05.475150

**Authors:** Sarah L. Fong, John A. Capra

## Abstract

**Motivation:** Thousands of human gene regulatory enhancers are composed of sequences with multiple evolutionary origins. These evolutionarily “complex” enhancers consist of older “core” sequences and younger “derived” sequences. However, the functional relationship between the sequences of different evolutionary origins within complex enhancers is poorly understood.

**Results:** We evaluated the function, selective pressures, and sequence variation across core and derived components of human complex enhancers. We find that both components are older than expected from the genomic background, and cores are enriched for derived sequences of similar evolutionary ages. Both components show strong evidence of biochemical activity in massively parallel report assays (MPRAs). However, core and derived sequences have distinct transcription factor (TF) binding preferences that are largely stable across evolutionary origins. Given these signatures of function, both core and derived sequences have substantial evidence of purifying selection. Nonetheless, derived sequences exhibit weaker purifying selection than adjacent cores. Derived sequences also tolerate more common genetic variation and are enriched compared to cores for eQTL associated with gene expression variability in human populations.

**Conclusions:** Both core and derived sequences have strong evidence of gene regulatory function, but derived sequences have distinct constraint profiles, TF binding preferences, and tolerance to variation compared with cores. We propose that the step-wise integration of younger derived and older core sequences has generated regulatory substrates with robust activity and the potential for functional variation. Our analyses demonstrate that synthesizing study of enhancer evolution and function can aid interpretation of regulatory sequence activity and functional variation across human populations.

**Significance Statement:** Thousands of human gene regulatory enhancers are mosaics of sequences from multiple evolutionary origins, yet how these different segments contribute to enhancer function is poorly understood. By dissecting their regulatory functions, transcription factor binding, constraint, and human genetic variation, we show that both older “core” and younger “derived” sequences in complex enhancers have strong evidence of gene regulatory function, but derived sequences are more likely to harbor genetic variants that influence function. Together, our results support a model in which the integration of sequences of different origins generates regulatory substrates with robust activity and the potential for functional variation.

## Introduction

Enhancers are distal, non-coding gene regulatory DNA sequences that modulate target gene expression in cell-type- and spatio-temporal-specific contexts (Shlyueva, Stampfel, and Stark 2014). Enhancer function is mediated by the binding of transcription factors (TFs) that recognize DNA sequence motifs and interact with promoters. Changes in enhancer function are major drivers of species divergence and variation within species (Wray 2007; Sholtis and Noonan 2010; Wittkopp and Kalay 2012; Franchini and Pollard 2015; Rebeiz and Tsiantis 2017), yet the evolutionary events underlying the creation and functional evolution of enhancers are less understood.

Enhancer activity turns over rapidly between mammalian species, but most sequences with current enhancer activity have ancient origins (Villar et al. 2015). Several connections have been discovered between the evolutionary origins of DNA sequences with enhancer activity and current gene regulatory functions. The age of a regulatory sequence is predictive of the genes that it likely targets, and different periods of regulatory sequence innovation have contributed to vertebrate evolution (Lowe et al. 2011). Moreover, younger mammalian neocortical enhancers are more weakly constrained, and many neocortical enhancers consist of sequences of multiple evolutionary origins (Emera et al. 2016). Underscoring the functional relevance of these evolutionary events, older sequences with gene regulatory activity are more enriched for heritability in a range of human complex traits than younger sequences with regulatory activity (Hujoel et al. 2019). These waves of regulatory change have been driven in large part by the integration of transposable elements (TEs) carrying different TF binding sites into the genome at different times (Marnetto et al. 2018).

Mammalian enhancer sequences are often composed of functional units, or modules, that bind different combinations of transcription factors (Long, Prescott, and Wysocka 2016; Jindal and Emma K. Farley 2021). Recent work has begun to reveal the nature of the modular organization of enhancer functions (Gotea et al. 2010; E. K. Farley et al. 2015; Tippens et al. 2020; Long, Osterwalder, et al. 2020; Wong et al. 2020). Enhancer sequences often result from the integration of different combinations of sequence over time (Emera et al. 2016; Fong and Capra 2021). However, models that synthesize the evolutionary origins of enhancer sequences with an understanding of functional modules are needed.

The potential value of integrating evolution and function is illustrated by existing models of protein-coding sequence evolution. Over evolutionary time, protein-coding sequences often generate novel protein functions by integrating functional modules in different combinations. Knowledge of the evolutionary origins of different proteins and domains provides valuable context for interpreting the evolution and function of protein families (Capra et al. 2013). As a result, many statistical frameworks exist for modeling protein domain and family evolution (Stolzer et al. 2015; Forslund, Kaduk, and Sonnhammer 2019). Expanding knowledge of the relationship between enhancer evolution and function will contribute valuable context for resolving gene regulatory functions of candidate disease variants of unknown significance, understanding the molecular basis for differences between species, and developing synthetic gene regulatory elements.

We recently explored how the evolutionary origins of an enhancer sequence are reflected in its functional and regulatory features, such as pleiotropy and robustness to perturbation of its biochemical activity by genetic variants (Fong and Capra 2021). We discovered that a significant fraction of enhancer sequences in diverse tissues consist of DNA from multiple evolutionary origins. These “complex” enhancers are the result of genomic integration and rearrangement events over evolutionary time. Complex enhancers are often active across multiple tissues, while their evolutionarily simpler counterparts are often active in tissue-specific contexts. Yet, the relationship between the sequences of different evolutionary origins in complex enhancers and the gene regulatory functions they produce is poorly understood. For example, whether all sequences from different evolutionary periods have independent gene regulatory functions is unclear in most complex enhancers.

Here, we address this gap by contrasting the evolutionary origins, functional characteristics, TF binding, selection pressures, and human genetic diversity of the oldest “core” regions and younger “derived” regions of complex enhancer sequences. We find that both core and derived regions have strong evidence of gene regulatory function, but derived regions have distinct properties in terms of their constraint profiles, TF binding preferences, and tolerance to variation compared with cores. In addition, complex enhancers show a strong enrichment for sequences of similar evolutionary ages. Overall, our results illustrate that the combination of core and derived regions in enhancer sequences often promotes robust gene regulatory activity while providing a substrate for functional variation in humans.

## Results

### Enhancers are commonly composed of older core and younger derived sequences

Thousands of human gene regulatory enhancers are composed of sequences with multiple evolutionary origins. Previous work separated the components of these “complex” enhancers into two classes—core and derived sequences (Figure 1A; Emera et al. 2016; Fong and Capra 2021). The “core” sequence(s) are the oldest sequences in an enhancer, and the younger sequence regions are “derived”. Our goal is to evaluate the function, selective pressures on, and sequence variation across these components of complex human gene regulatory enhancers genome-wide by studying the ages of enhancer sequences (Figure 1A).

**Figure 1:**
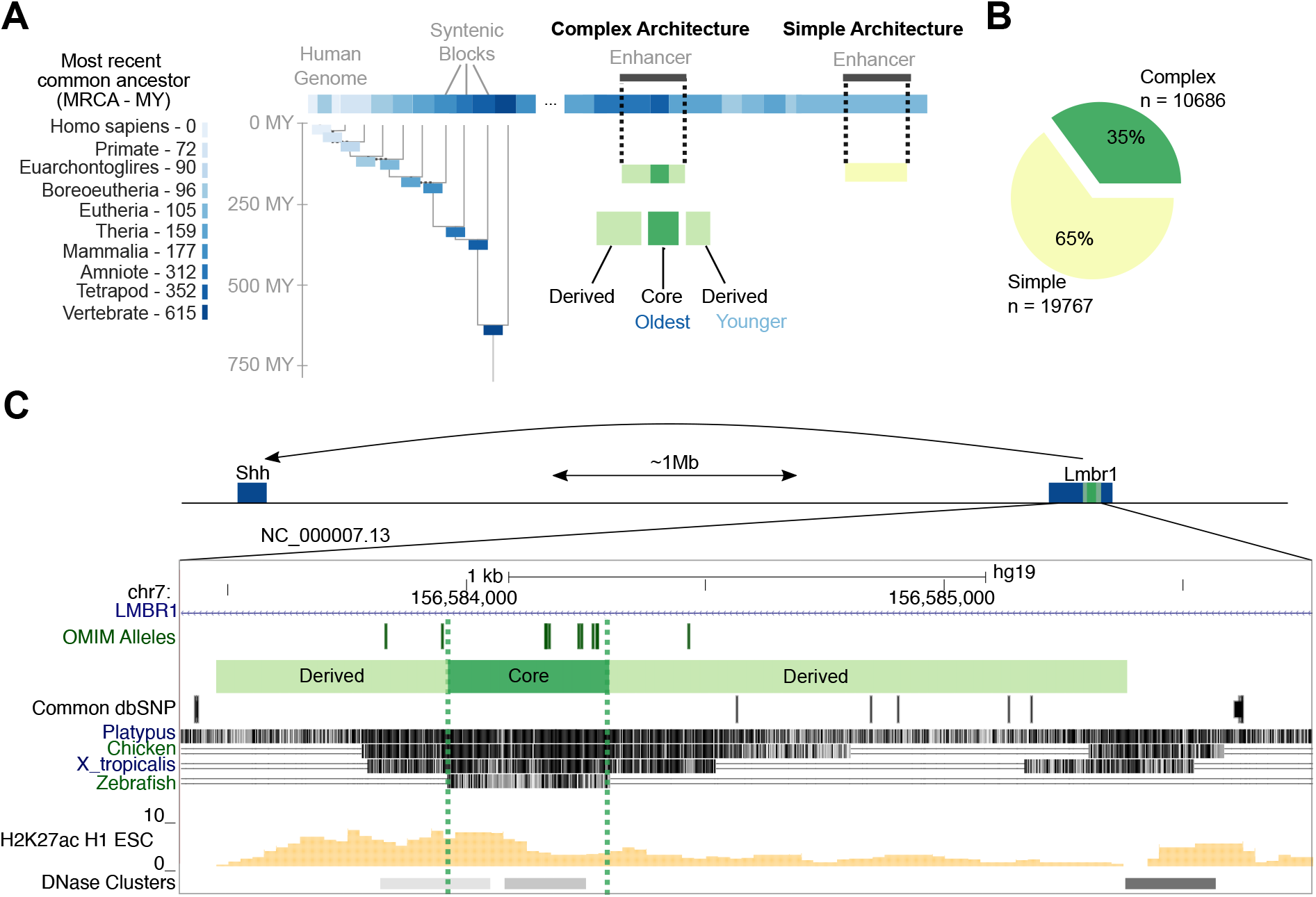
Complex enhancers consist of older core and younger derived sequences. **(A)** Illustration of the approach for mapping enhancer sequence ages and architectures. We quantify the age of a sequence with human enhancer activity based on the oldest most recent common ancestor (MRCA) in overlapping syntenic blocks from the MultiZ multiple sequence alignments of 46 vertebrates (color key). Enhancer age is assigned as the oldest, overlapping syntenic block age. Estimates of divergence time in millions of years ago (MY) from TimeTree (Hedges et al., 2015) are annotated in the color key. **(B)** Autosomal transcribed enhancers from the FANTOM5 consortium (N = 30,434) were classified as having complex (multi-age) or simple (single-age) architectures. Complex enhancers were further dissected into the oldest “core” and younger “derived” sequence regions. **(C)** A complex developing limb bud enhancer (NC 000007.13) of *SSH* is located ∼1 Mb away in an intron of *LMBR1* and has multiple evolutionary origins. Among 11 variants in OMIM that cause preaxial polydactyly 2 (PPD2), eight variants are in the Vertebrate core region, and three are in the Tetrapod derived region. Common variants (minor allele frequency *>*1% in 1000 Genomes Project phase 3) from dbSNP (version 153) are observed only in derived regions. H3K27ac ChIP-seq peaks in H1-ESC and DNase I hypersensitive clusters from 125 cell lines in ENCODE3 are shown for context. age 0.175 observed v. 0.152 expected; MWU p *<* 2.2e-238). This indicates that both core and derived sequences are older than expected and suggests that both components often have constrained regulatory function.

To illustrate the components of a complex enhancer, we dissected evolutionary origins of the zone of polarizing activity regulatory sequence (ZRS), a long-range enhancer of *SHH* involved in developmental limb bud formation (Laura A. Lettice, Devenney, et al. 2017). The ZRS sequence achieves its regulatory function via the consolidation of multiple distinct regulatory domains (L. A. Lettice 2003, Laura A. Lettice, Williamson, et al. 2012, Long, Prescott, and Wysocka 2016). The core sequence has origins before the last common ancestor of all vertebrates, and it is flanked on both sides by multiple derived regions with origins in the ancestors of tetrapods, amniotes, and mammals. This enhancer sequence is both strongly conserved and involved in evolutionary variation in limb morphology. Loss of function variants at this locus contributed to limbless evolution in snakes (Kvon et al. 2016), while variants in vertebrate and tetrapod sequences are associated with preaxial polydactyly 2 (PPD2) (Hill and Laura A. Lettice 2013; Ushiki et al. 2021). In humans, eight of the eleven PPD2-causing variants annotated in the Online Mendelian Inheritance in Man (OMIM) catalog are located in the Vertebrate core of the ZRS enhancer sequence, while three are located in Tetrapod derived regions (Figure 1C). Common variants (minor allele frequency *>* 1% in 1000 Genomes Projects from dbSNPv153) are observed in the younger derived amniote and mammal sequences, but not in older tetrapod and vertebrate sequences. This example demonstrates that although disease variants are concentrated in the core sequence, point mutations in both older core sequences and younger derived regions can cause human disease.

### Derived regions constitute a substantial fraction of complex enhancer sequences

We first evaluated the basic features of core and derived sequences in a set of complex, non-coding autosomal transcribed enhancers active in 112 diverse tissues and cell samples from the FANTOM5 consortium (N = 10,686; Figure 1B). Derived regions represent 46% of the base pairs (bp) in a typical complex enhancer sequence (median total length of 310 bp) (Figure 2A, left), and complex enhancers have a median of one derived region per core region (Figure S1). However, derived regions are shorter than core regions (Figure 2A, right; median bp 136 derived v. 174 core) and shorter than expected from sets of shuffled length- and chromosome-matched non-coding genomic background regions with multiple sequence ages, which we refer to as “expected” (Figure S2; median bp 136 observed v. 157 expected; Mann-Whitney U (MWU) p = 1.4e-46). Conversely, core regions are longer than expected (median bp 174 observed v. 143 expected; MWU p = 2.4e-73; Figure S2). Stratifying enhancers by their core ages and repeating these comparisons yielded similar results (Figure S3). Thus, derived sequences make up a substantial fraction of complex enhancer sequence and are sufficiently long to bind combinations of TFs.

**Figure 2:**
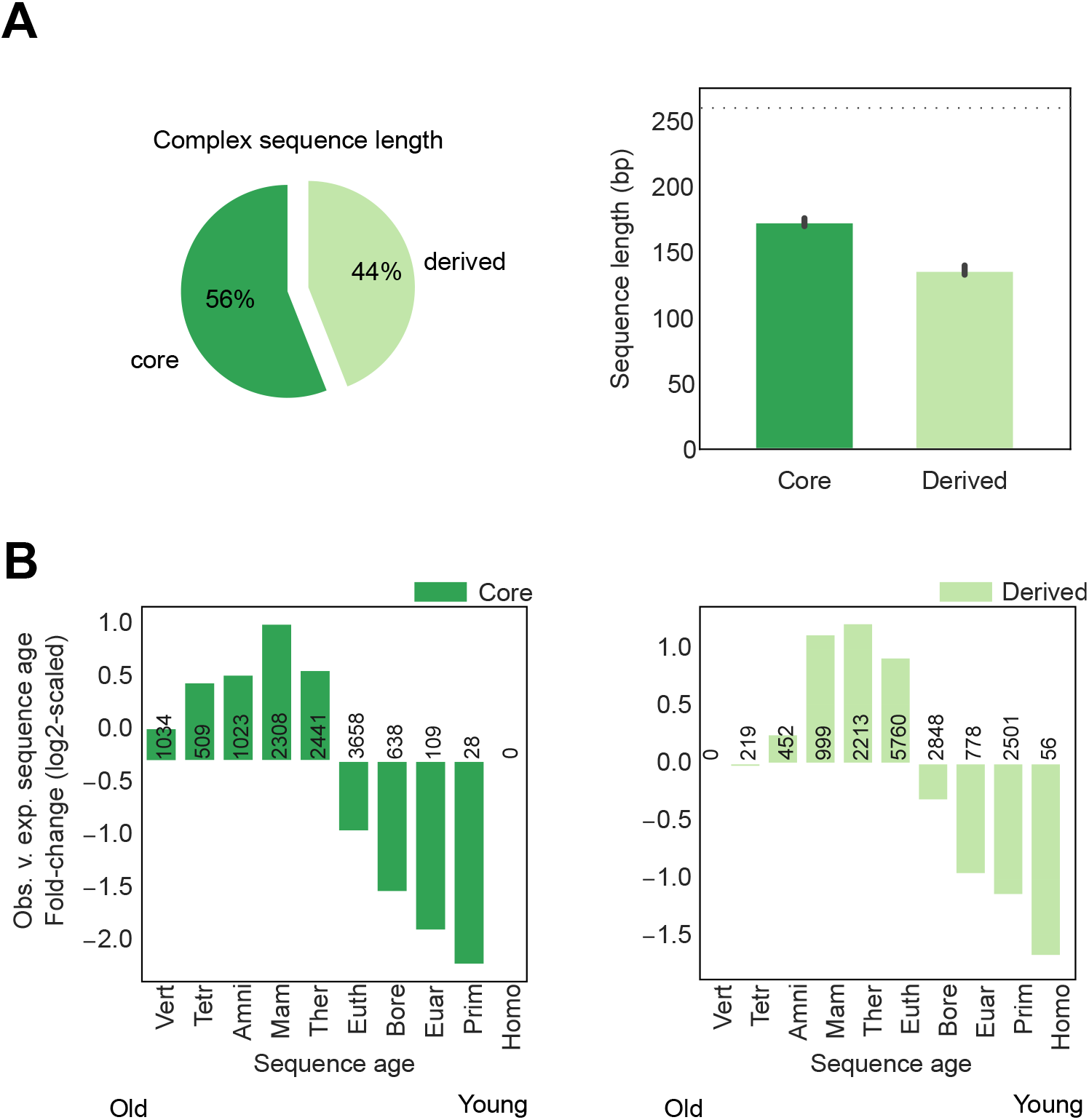
Derived sequences are shorter than cores and older than expected from the non-coding genome. (**A**) Derived regions constitute 44% of complex enhancer sequences (left), but are shorter than core regions (right, median 136 bp derived v. 174 bp core, Mann Whitney U p = 4.9e-57). Both core and derived regions are shorter than simple enhancers (dashed line, median 260 bp simple, p *<* 2.2e-308). (**B**) Both core and derived sequences are enriched for older sequence ages and depleted of younger sequence ages. Per age, the log_2_ of the fold change of the observed core (left) and derived (right) sequence ages versus the expected proportion estimated from 100x sets of length-, chromosome-, and architecture-matched shuffled non-coding sequences. Sample size is annotated per bar.

### Both derived and core regions are older than expected from the non-coding genome

Enhancer sequences are older than expected from the non-coding genomic background, suggesting that they have likely been maintained due to their function (Lowe et al. 2011; Villar et al. 2015; Emera et al. 2016; Marnetto et al. 2018; The ENCODE Project Consortium et al. 2020; Fong and Capra 2021). We expanded previous analyses of enhancer ages to consider the multiple evolutionary origins of complex enhancers. We compared the distributions of core and derived sequence ages to expected sequences with matched core ages. Core sequences are enriched for older ages (Therian ancestor and older) compared with expected core sequence ages (Figure 2B left; median age 0.30 observed v. 0.175 expected; MWU p *<* 2.2e-238). Derived sequences are also enriched for older ages compared to derived regions of expected sequences, and the enrichment extends to sequences with Eutherian origins (Figure 2B right; median derived sequence

### Complex enhancers are enriched for core and derived sequences from consecutive phylogenetic branches

To explore whether core and derived sequences in the same complex enhancer have temporal relationships, we evaluated enrichment for sequence age combinations among observed derived and core sequence pairs. We hypothesized that derived sequence origins would likely occur soon after the origins of the corresponding core sequences. Overall, enhancers are enriched for core and derived sequences from the consecutive phylogenetic branches compared to expected complex regions from random, length- and chromosome-matched, non-coding genomic background with multiple sequence ages (Figure 3). This suggests a preference for integration of derived sequences into older core enhancer sequences on contiguous branches, and that integration of much younger derived sequences was less tolerated by old cores. In addition, Mammalian core sequences and older are enriched for Therian derived sequences and older, but depleted of derived sequences from younger ages. The oldest complex enhancers (from the Mammalian ancestor and earlier) are enriched for derived sequences of several ancient origins (from the Therian ancestor and earlier), likely due to their very old ages. These results indicate that the pairing of core and derived sequences within complex enhancers is not random with respect to their origins and that evolution favors the step-wise addition of derived sequences that are near in age to the core sequence.

**Figure 3:**
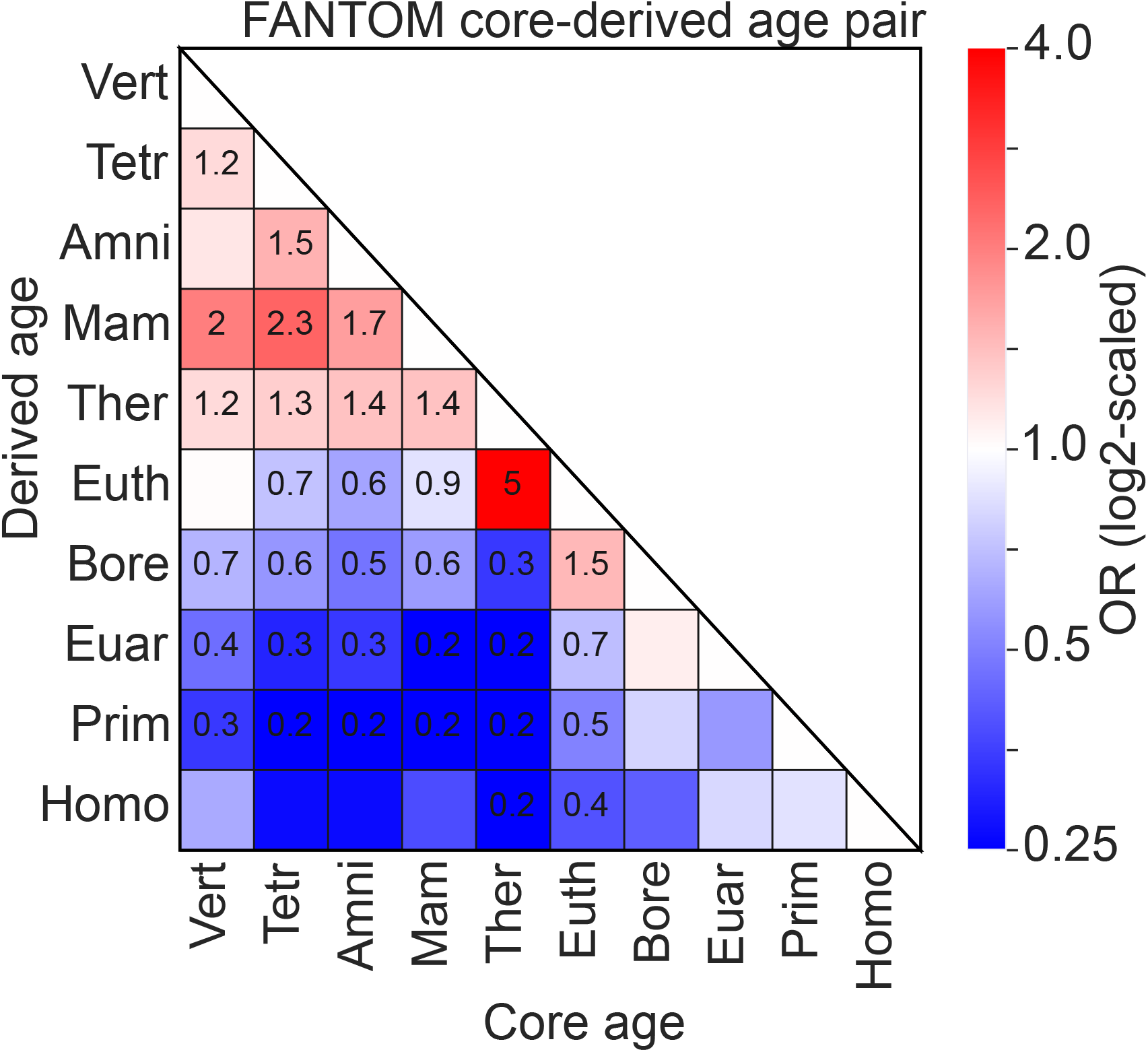
Complex enhancers are enriched for core and derived sequences from consecutive phylogenetic branches. For each enhancer core age, the enrichment for derived sequences of each age was measured against the expectation from core-age-matched shuffled sequences using Fisher’s exact test. The boxes are colored according to the *log*_2_ of the corresponding odds ratios (OR). Text in a box indicates significant enrichment (red) or depletion (blue) after controlling the false discovery rate at 0.05 with the Benjamini-Hochberg procedure. The oldest complex enhancers (pre-placental mammals) are enriched for older derived sequences. Outside of the oldest enhancers, there is consistent significant depletion for complex enhancers with core and derived segments with origins on non-consecutive phylogenetic branches.

### Derived sequences have higher transcription factor binding site density than cores

Transcription factor (TF) binding at enhancer sequences is required for gene regulation, but the relative contributions of core and derived sequences to TF recruitment in complex enhancer sequences is not known. Some derived regions may be non-functional sequences flanking functional enhancer cores that are identified for technical reasons. Alternatively, derived sequences may bind TFs essential for the proper regulation of gene expression in specific contexts.

To evaluate the role of derived sequences in binding TFs, we leveraged the ENCODE project’s deep characterization of TF binding sites and enhancers in HepG2 and K562 cells: 119 and 249 TF chromatin immuno-precipitation sequencing (ChIP-seq) assays and previously identified candidate cis-regulatory elements (cCREs) with enhancer-like signatures based on DNase I hypersensitivity, CTCF, and histone mark ChIP-seq assays (The ENCODE Project Consortium et al. 2020). We first confirmed that our findings on complex HepG2 and K562 enhancer architectures are consistent with those in FANTOM5 (Figure S5, Figure S8). We then quantified TF binding site (TFBS density and enrichment patterns in core and derived regions of these enhancers. In complex HepG2 enhancers, we observe that 46% of derived regions bind TFs compared to 67% of core regions and 87% of simple HepG2 enhancers (Figure S17). A similar trend was observed in K562 complex enhancers, where 59% of derived, 79% of core, and 93% of simple regions bind TFs. We note that we have better power to detect TFBS in K562 cells because more ChIP-seq assays have been run in that cell model (249 K562 v. 119 HepG2 ChIP-seq assays). Complex enhancer regions with no evidence of TF binding occur at similar frequencies across ages for both HepG2 and K562 cells, suggesting that TF binding evidence is independent of enhancer sequence age (Figure S6).

In complex HepG2 enhancers with bound TFs, derived regions have higher TFBS densities compared to core regions and simple enhancers (Figure 4A; median 4.3 binding sites/100 bp in derived regions versus 3.6 binding sites/100 bp in core regions, MWU p = 1.1e-68). We observed a similar trend in complex K562 enhancers (Figure S9A; median 7.4 binding sites/100 bp in derived regions versus 6.4 binding sites/100 bp in core regions, MWU p = 3.5e-52). This trend of higher derived region TFBS density is consistent across enhancers of different ages (Figure S7), suggesting that derived sequences bind TFs and have higher TFBS densities than core sequences across evolutionary ages. Thus, derived sequences have a higher density of assayed TF binding sites when a binding site is present, but they are less likely to be bound by a TF than core segments overall.

**Figure 4:**
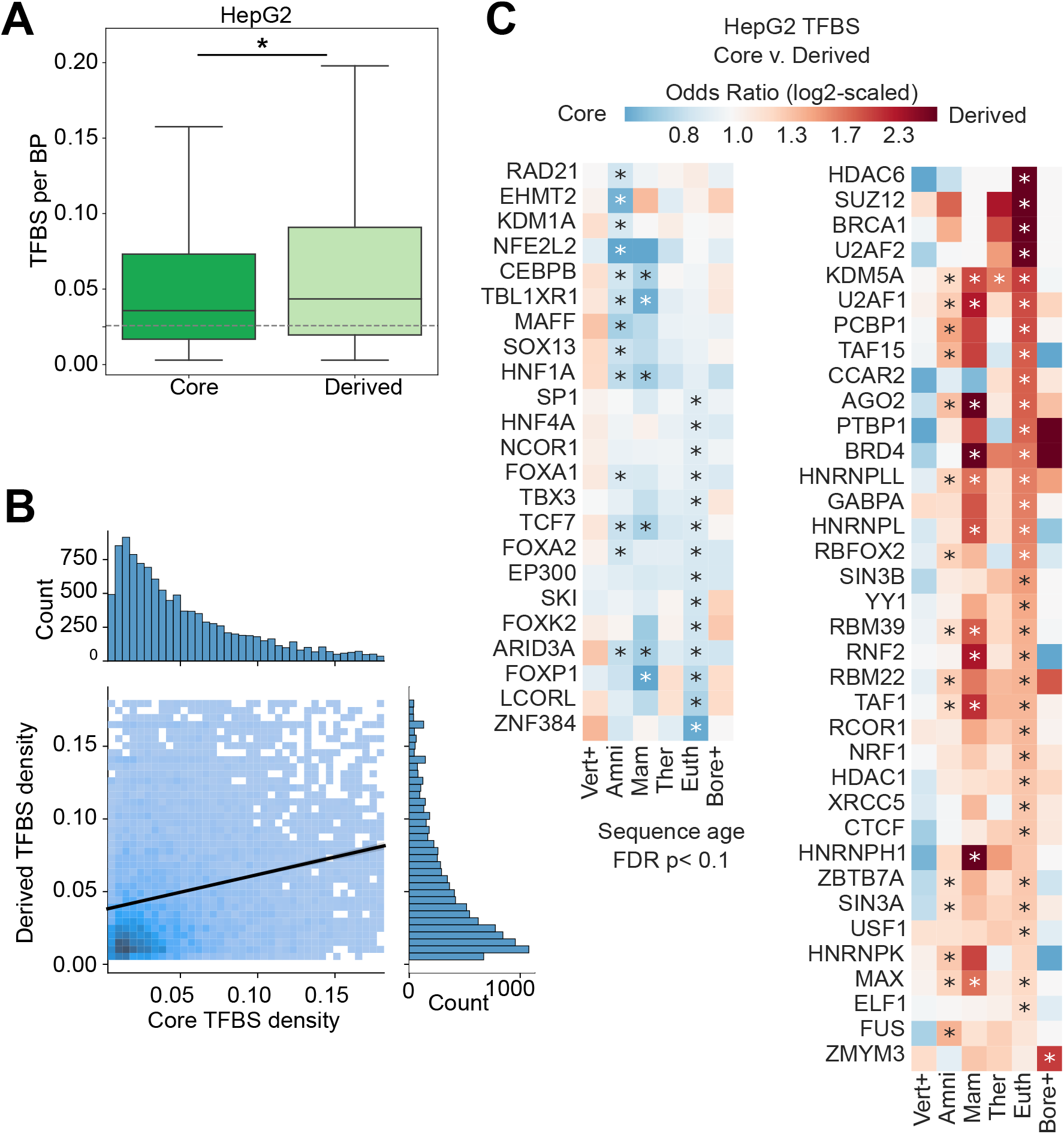
Derived regions have high transcription factor binding site densities and bind different transcription factors compared to core regions. (**A**) Among complex enhancers with evidence of TF binding in either core or derived regions (N = 20263), derived regions (N = 20210) have higher TFBS densities than core regions (N = 19957) (mean 0.043 derived v. 0.036 core TFBS/bp; Mann Whitney U p = 1.1e-68). However, derived regions are more likely to have no TFs bound than core regions (Figure S17). Core and derived regions both have higher TFBS density than simple enhancers (dashed line; 0.026 TFBS/bp). **(B)** TFBS density is positively correlated between core-derived sequence pairs within complex enhancers with evidence of TF binding in both core and derived regions (N = 11899). Color intensity represents the density of core-derived pairs, and the black line is a linear regression fit (slope=0.23, intercept=0.04, r=0.24, p = 5.1e-140); outliers (*>*95th percentile) are not plotted for ease of visualization. **(C)** Derived and core regions of the same age are enriched for binding of different TFs and enrichment patterns are generally consistent across ages. TFBS enrichment for each age was tested using Fisher’s exact test; only TFs with at least one significant enrichment (FDR *<* 0.1) are shown. Vertebrate, Sarcopterygii, and Tetrapod enhancer ancestors were grouped into “Vert+”. Boreotherian, Euarchontoglires, and Primate enhancer ancestors were grouped into “Bore+”.

Next, we quantified the relationship of TFBS density within core and derived segments of the same complex enhancer. Among HepG2 enhancer sequences with bound TFs in both core and derived sequences (N = 11899), TFBS density is positively correlated between the core and derived regions (Figure 4B; linear regression slope=0.23, intercept=0.04, r=0.24, p=5.1e-140). We observed a similar positive correlation in K562 cells (Figure S9; linear regression slope=0.39, intercept=0.056, r=0.39, p = 0.0, stderr=0.008). Relaxing our criteria to include core and derived sequences with no evidence of TF binding, we still observe that core and derived density within a single enhancer sequence is positively correlated (Figure S10). These results show that TFBS density is overall positively correlated in adjacent core and derived regions, and that when bound, derived sequences have a higher TFBS density.

### Core and derived sequences are enriched for distinct TFBS across ages

Given the differences in TF binding probability and density between core and derived regions, we hypothesized that regions might also exhibit different TF preferences. Indeed, we found that derived and core HepG2 enhancer regions are enriched for binding of distinct TFs (Figure 4C). Furthermore, many TFs are consistently enriched in derived or core regions across multiple sequence ages, suggesting that specific TFs have a preference for binding core or derived sequence contexts. GO annotation enrichment analyses did not identify strong specific functional enrichment among TFs with different binding preferences. No GO annotations were enriched in derived sequences at any age. However, core sequence TFs with preferences for the Amniota and Eutherian ancestors are enriched for “regulation of transcription by RNA polymerase II” (GO:0006357, derived v. core odds ratio (OR) = 0.13, p = 0.03 for Eutherian and OR = 0.08 p = 0.04 for Amniota sequences, FDR *<* 10%). This suggests that core TFs are enriched for factors that recruit the RNA polymerase II machinery needed to initiate transcription, while derived TFs are depleted and may instead diversify transcriptional activity. We tested these conclusions in another deeply characterized ENCODE cell line, K562, and found similar patterns (Figure S8), including higher TFBS density in derived sequences and TF-DNA binding biases in core and derived sequences (Figure S9). TFs specific to core and derived sequences were unique among HepG2 and K562 enhancers, suggesting that core and derived sequence evolution is cell-type-specific. Overall, these results indicate that many derived regions have distinct TF binding partners from their associated cores.

### Core and derived regions have similar activity in MPRAs

Given that TF binding in core and derived sequences, we hypothesized that these regions often have functional gene regulatory activity. To evaluate this, we compared the estimated activity of core and derived enhancer sequences from previously published SHARPR massively parallel reporter assays (MPRAs) (Ernst et al. 2016. Briefly, SHARPR uses probabilistic graphical models to estimate base-pair-level biochemical activity from the levels of transcribed mRNA and corresponding episomal DNA plasmids for 4,000 HepG2 and K562 enhancers. We assigned ages and architectures to the sequences with per bp regulatory activity in SHARPR-MPRA assays (*>*1:1 ratio of mRNA transcripts to DNA plasmids). Among active bases, Derived and core sequences have similar activity per bp in both K562 and HepG2 cells, though core regions are slightly higher (Figure 5; HepG2: median per bp activity 1.58 derived v. 1.65 core, MWU p = 2.0e-6; K562: 1.50 derived v. 1.63 core, p = 6.6e-32). Stratified by age, we do not observe any consistent trends in core v. derived activity across evolutionary periods in HepG2 or K562 cells (Figure S12). Simple enhancers (i.e., enhancers of a single age) show slightly higher activity per bp (median 1.69) than both core and derived segments of complex enhancers. Nonetheless, these data suggest that many derived sequences are biochemically active, have similar levels of activity compared with their adjacent cores, and contribute to gene regulatory function.

**Figure 5:**
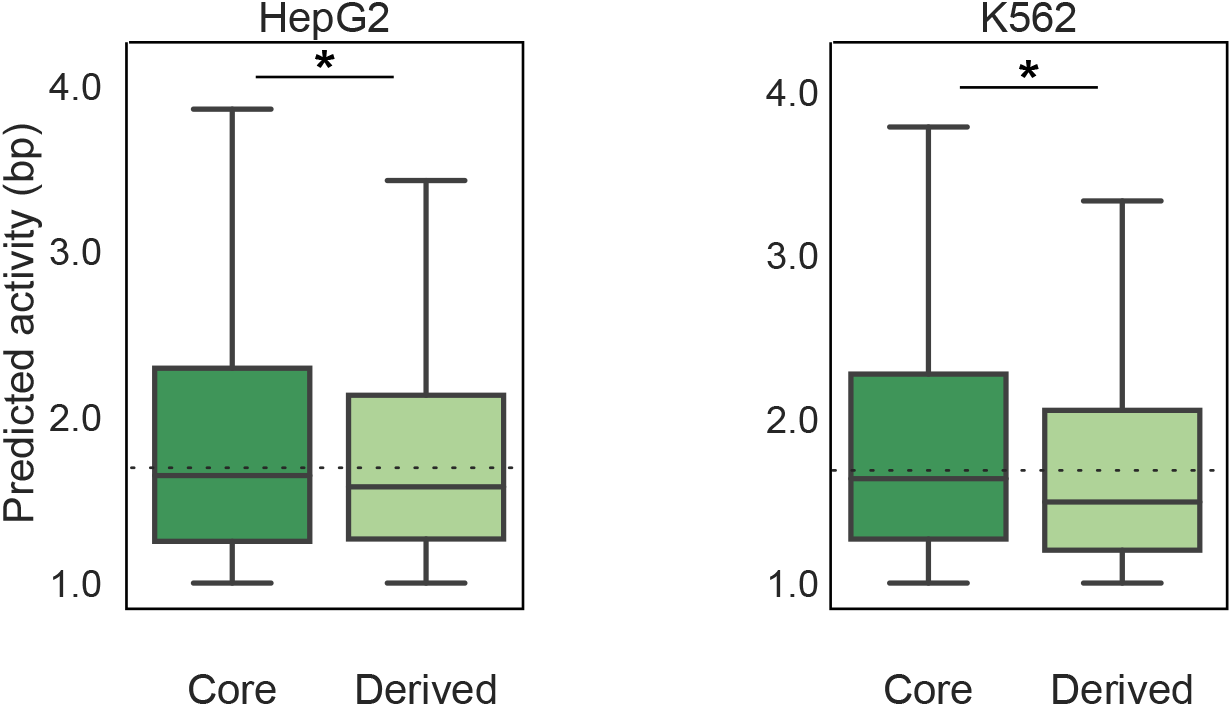
Both core and derived regions have regulatory activity in massively parallel reporter assays. Many derived sequences had enhancer activity in a previous HepG2 MPRA analysis (activity score 1 for entire enhancer; N = 2000 enhancers for HepG2 and N = 2000 for K562; Ernst et al. 2016). However, derived activity was modestly, but significantly lower than core sequences (N = 9076 bps tested) (median 1.58 derived v. 1.65 core activity per bp; Mann Whitney U p = 2.0e-6) Patterns were similar in K562 cells (mean 1.50 derived v. 1.64 core; p = 6.6e-32). Both core and derived segments of complex enhancers had lower activity per base pair than simple enhancers (dashed lines, median 1.69). For HepG2, N = 6498 derived and N = 9076 core active bp were tested, while for K562, N= 7568 derived and N = 9846 core bp were tested.

### Derived sequences are less evolutionarily constrained than core sequences

We next evaluated evolutionary constraints on core and derived sequences. To do this, we compared LINSIGHT per bp estimates of purifying selection (Huang, Gulko, and Siepel 2017) for derived sequences and associated cores in the FANTOM dataset. Overall, derived sequences have slightly, but significantly lower LINSIGHT scores than adjacent cores (Figure 6A; median 0.07 derived v. 0.08 core LINSIGHT score; derived v. core MWU p ¡2.2e-238), suggesting that derived regions experience weaker purifying selection than adjacent enhancer cores. This pattern also holds when stratifying complex enhancers by sequence age (Figure S14). As older enhancer sequences are generally under stronger evolutionary constraint, we also compared core and derived sequences of the same age and found that derived regions also have consistently lower LINSIGHT scores than age-matched core sequences (Figure S13). Together, these results indicate that derived sequences are under slightly weaker purifying selection than neighboring core regions in the same complex enhancer and than core regions of the same age.

**Figure 6:**
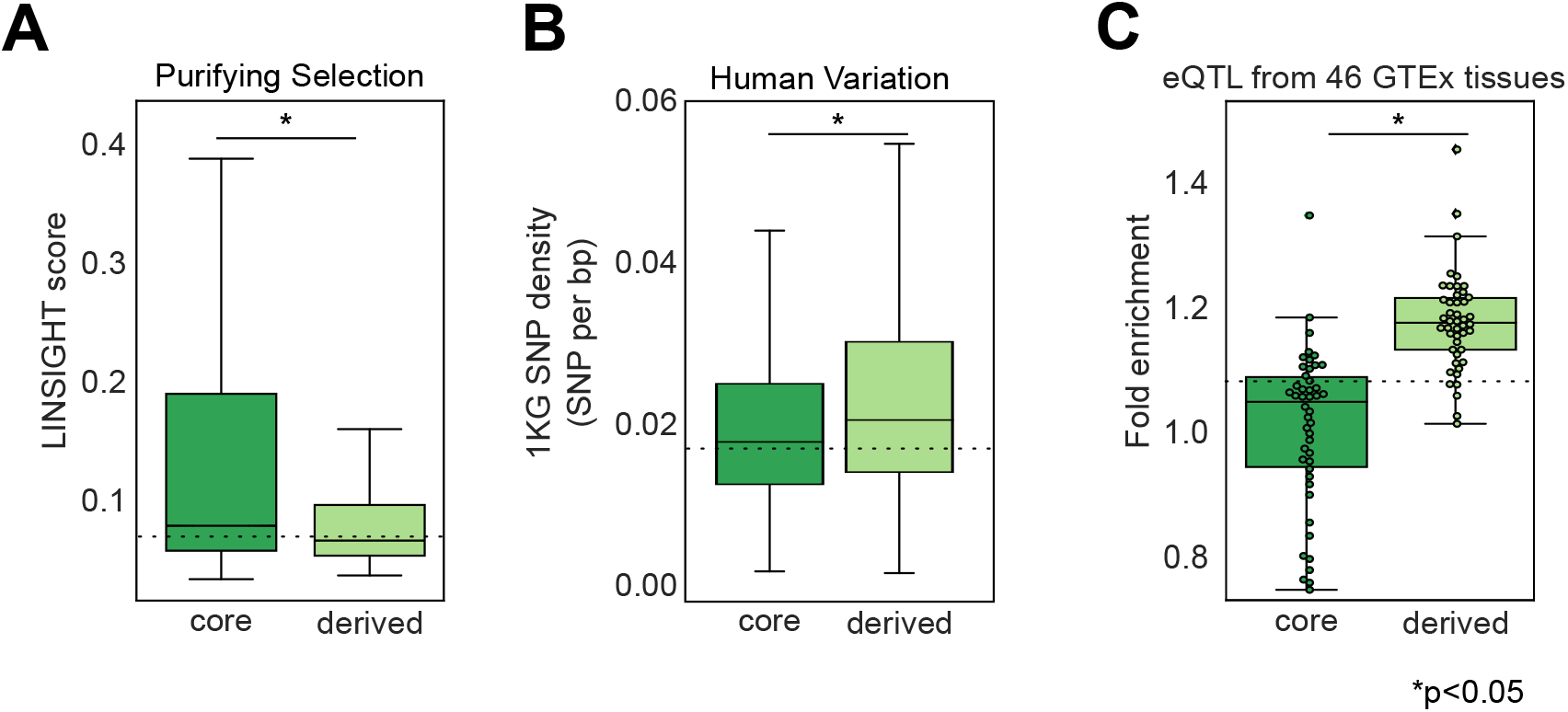
Derived regions experience weaker purifying selection, have more genetic variation, and are enriched for eQTL compared to adjacent core sequences. (**A**) Derived regions (N = 2,021,098 bp) have significantly lower LINSIGHT purifying selection scores than adjacent core regions (N = 2,271,279 bp; median 0.08 core v. 0.07 derived per bp LINSIGHT score MWU p *<*2.2e-308). The dashed line represents median simple enhancer LINSIGHT score (0.07, N = 5,398,405 bp). **(B)** Derived regions (N = 26,451 variants) have higher genetic variant densities than associated core regions (N = 27,691 SNPs; median 0.020 derived v. 0.018 core SNPs per bp; MWU p = 1.4e-202). Variant densities were calculated as the number of variants from the 1000 Genomes Project in each enhancer region divided by its length. The dashed line represents median density in simple enhancers (N = 71,415 SNPs). **(C)** Derived regions are significantly more enriched for expression quantitative trait loci (eQTL) than core regions (Kruskal-Wallis p = 9.4e-12). eQTL from the GTEx consortium v6 from 46 tissues were intersected with enhancers. Enrichment for eQTL from each tissue in core and derived components was estimated from 1000 length-matched, chromosome-matched permutations, and confidence intervals were estimated from 10,000 bootstraps. Each dot represents enrichment for eQTL of a tissue in core or derived enhancer regions. Derived, core, and simple enhancers have significantly different eQTL enrichments (MWU p = 1.3e-11). The dashed line represents the median simple enhancer eQTL enrichment across 46 tissues.

### Derived enhancer regions have more genetic variation than core regions

Given the modest differences in purifying selection between core and derived sequences, we compared their variant densities using genetic variants segregating in diverse human populations from the 1000 Genomes Project. As expected, derived sequences have modestly higher variant densities than complex core regions (Figure 6B; median 0.020 v. 0.018 variants per bp; MWU p = 1.4e-202) and than simple enhancers (median 0.017 variants per bp). Consistent with this, global minor allele frequencies are also slightly higher in derived sequences compared to core and simple sequences (Figure S15). This implies that derived sequences accumulate more genetic variants than core sequences, consistent with our observation that derived regions are under weaker purifying selection than adjacent cores.

### Derived enhancer regions are enriched for eQTL

To explore whether variation in derived regions is associated with changes in their effects on gene regulation, we quantified enrichment of expression quantitative trait loci (eQTL) in derived and core regions using eQTL from GTEx for 46 tissues (GTEx Consortium 2017). As expected, all enhancer architecture components are enriched for eQTL compared to the genomic background (Figure 6C; median OR 1.20 derived, 1.50 core, 1.10 simple; MWU core v. derived, p = 1.3e-11). However, derived regions have the strongest enrichment. This is consistent with the higher minor allele frequencies (Figure S15) and lower purifying selection pressure (Figure 6A) in derived regions. Nonetheless, eQTL enrichment in derived sequences indicates that variation in these regions of complex enhancers contributes to gene expression variability in human populations.

## Discussion

Our dissection of human transcribed enhancers reveals that a substantial fraction (∼35%) is composed of sequences that originate from multiple evolutionary periods. We demonstrate that both the older core and younger derived sequences in these evolutionarily complex enhancers often show evidence of biochemical function and evolutionary constraint. Complex enhancers are enriched for core and derived sequences of similar ages. This suggests that the evolution of complex enhancer sequences proceeded in a step-wise and temporally-constrained manner. However, we observe important differences in core v. derived regions, including the density and identity of TFs that bind, evolutionary constraint, and genetic variation in these different sequence contexts. We confirm previous results from neocortical enhancers that derived regions are generally under less constraint Emera et al. 2016. We also find that they are more likely to harbor genetic variation in human populations and variants that are associated with gene expression levels. Thus, both core and derived sequences appear to often be functional, but they also exhibit different evolutionary and functional attributes.

These results motivate further investigation of how the evolutionary origins of enhancer sequences relate to their functions and suggest a potential analogy to protein function and evolution. Protein sequence functional architecture is intimately tied to evolution; most proteins are composed of modular domains with independent evolutionary origins that are juxtaposed in different functional combinations. However, many fundamental questions remain to be resolved about the modularity of enhancer evolution and function:

### What is the functional importance of derived enhancer sequences to their core regions?

Our results suggest that core and derived sequences often both have gene regulatory functions. However, many questions about the sufficiency of core and derived sequences remain to be addressed. For example, we do not know how often core and derived sequences have stand-alone regulatory activity. Previous work has proposed that promoters and enhancers have many similar features, including transcription start sites, bidirectional transcription, and GC-rich sequences (Andersson and Sandelin 2020), even though promoters require enhancer sequences to increase gene expression. Derived regions have slightly higher GC content than cores (Figure S**??**), have higher activity, and are less evolutionarily conserved than core sequences. Thus, it is possible that derived regions may function to enhance the promoter-like activity of core enhancer regions. In other words, derived sequences may enhance core enhancer activity.

Future studies should assess how often core enhancer sequences are sufficient for gene regulatory activity without flanking derived regions, and when core and derived regions cooperate to specify regulatory function. We anticipate that both scenarios may be common among complex enhancers. Further, the molecular mechanisms by which the core and derived regions contribute to regulatory function (e.g., changing chromatin accessibility, binding different TFs) must be determined. Many of these questions can be answered with evolution-aware reporter assays and gene editing strategies that disrupt core or derived sequences while preserving other sequence properties.

### Are evolutionary modules functional modules?

Functional dissection of enhancer sequences suggests the modular organization of many enhancers (Dukler et al. 2017; Long, Prescott, and Wysocka 2016; Sabarís et al. 2019). Previous work has focused on this modularity in the context of TFs and other functional genomic markers. These have revealed the importance of transcriptional units (Tippens et al. 2020), the organization of its TFBS into clusters (Gotea et al. 2010), and the spatial distribution between TFBS (E. K. Farley et al. 2015; Grossman et al. 2018) to enhancer sequence modularity. Taking an evolutionary perspective, we demonstrate that many enhancers consist of distinct evolutionary modules. Yet, how these evolutionary modules relate to functional modules must be further clarified. For example, different evolutionary modules could have distinct modular regulatory functions that are combined, as is common in proteins. The independent biochemical activity for many derived enhancer sequences suggests that this scenario occurs. Further, core and derived sequences may develop synergistic regulatory functions. A recent analysis of *SOX9* gene regulation has demonstrated that two sub-regions of the EC1.45 enhancer (from Therian and Vertebrate common ancestors, respectively) synergistically activate human *SOX9* expression(Long, Osterwalder, et al. 2020). The extent to which synergy is observed between core and derived regions of complex enhancer sequences should be explored further. We speculate that the combination of sequences from different evolutionary origins often enables gene regulatory innovation while conserving core regulatory functions. As suggested in the previous section, future work should combine evolutionary analysis with high-resolution assays of regulatory function to assess the relationship between evolutionary sequence modules and function.

### Can considering enhancer evolutionary architecture aid interpretation of rare and common genetic non-coding variation?

Our work suggests that considering the evolutionary history of core and derived regions may provide valuable context for interpreting the function and disease relevance of human variation. The *SHH* enhancer provides an example where rare variants causing PPD2 are more prevalent in the core region and common variants are only present in the derived segments. Whether deleterious rare variation is generally concentrated in enhancer cores must be explored further. However, the small number of known non-coding Mendelian variants makes enrichment analyses challenging. With regard to common variation and associations with complex traits, we observed that eQTL are enriched in derived sequences. Derived regions also have higher variant density and slightly higher minor allele frequency than core regions; thus, we have greater power to detect effects on gene expression. Given the presence of linkage disequilibrium, whether variants in derived sequences directly affect gene expression variation must be tested to estimate their true contribution. Recent work has reported that the heritability of common variants is overrepresented in older gene regulatory elements (Hujoel et al. 2019), but whether this signal is due to variation in older complex enhancers and more specifically in cores, derived regions, or both remains to be explored. In general, more work is needed to understand the implications of common and rare variation in enhancer cores, derived regions, and their association with human traits.

### Limitations

Our work has several limitations. The available sequence, TF, and functional data limit the scope and resolution of some analyses. First, the sampling of species with available genome sequences, the depth of sequencing, and the quality of available genome assemblies all influence estimates of sequence age (Sholtis and Noonan 2010; Margulies and Birney 2008). Moreover, varying levels of constraint over time also influence sequence age estimates. Thus, the age estimates should be considered a lower bound. Moreover, we emphasize that the estimated age of sequences with human enhancer activity is not necessarily the age when the sequence first gained enhancer activity. Second, the TF binding site analyses focus on the 30 bp centers of ChIP-seq peaks. Given the variation in peak shapes and sizes of regulatory motifs, our heuristic may not always capture the exact position of TF binding. Nonetheless, given the large number of TFs and enhancers considered, we anticipate that the observed trends hold in general. Third, we leveraged previously published MPRA data; however, these only covered a few thousand enhancer regions in two cellular contexts. Without further biochemical assays, we cannot test whether most core and derived sequences have regulatory activity when separated. This is an important avenue for future work to determine whether derived sequences enhance pre-existing enhancer activity or if they work with core sequences to nucleate enhancer activity. Fourth, due to the challenges of linking regulatory elements to genes, we do not evaluate the gene targets associated with complex enhancers. Given their age and persistence over long evolutionary time, we speculate that complex enhancers often regulate genes involved in essential processes (Berthelot et al. 2018). Finally, in the TFBS analyses, we do not explore the properties of enhancers lacking TFBS in core or derived regions (N = 12085/27548, or 43% of HepG2 and N = 8545/22687 or 38% of K562 complex enhancers with no derived TFBS overlap. These may be misclassified simple enhancers (i.e., enhancer sequences for which the functional sequence is of a single age) or they may bind TFs or spatial combinations of TFs that have not been previously characterized. Given that these represent a small fraction of all enhancers and that they are likely enriched for false positives, we do not anticipate that this should influence our main conclusions.

### Conclusion

Variation in gene regulatory sequences underlies much of the phenotypic variation between individuals and species. However, unlike protein sequences, we do not understand how enhancer sequence origin and evolution relate to functional activity. Here, we show that enhancers commonly consist of sequences from multiple evolutionary epochs and that both core and derived segments regularly exhibit hallmarks of gene regulatory function. Thus, our results support and extend previous models of modular enhancer evolution by sequence accretion (Emera et al. 2016, Fong and Capra 2021)and suggest that enhancer sequences composed of evolutionary units may promote gene regulatory function and variability in gene expression. Our work motivates the further study of the evolution of gene regulatory elements and the functional interaction of sequences of different origins over evolutionary time.

## Methods

### Assigning ages to sequences based on alignment syntenic blocks

The genome-wide hg19 46-way and hg38 100-way vertebrate multiz multiple species alignment was downloaded from the UCSC genome browser. Each syntenic block was assigned an age based on the most recent common ancestor (MRCA) of the species present in the alignment block in the UCSC all species tree model (Figure 1A). For most analyses, we focus on the MRCA-based age, but when a continuous estimate is needed we use evolutionary distances from humans to the MRCA node in the fixed 46-way or 100-way neutral species phylogenetic tree. Estimates of the divergence times of species pairs in millions of years ago (MYA) were downloaded from TimeTree (Hedges et al. 2015). Sequence age provides a lowerbound on the evolutionary age of the sequence block. Sequence ages could be estimated for 93% of the autosomal base pairs (bp) in the hg19 human genome and 94% of the autosomal bp in the hg38 human genome.

### eRNA enhancer data, age assignment, and architecture mapping

We considered enhancer RNAs (eRNAs) identified across 112 tissues and cell lines by high-resolution cap analysis of gene expression sequencing (CAGE-seq) carried out by the FANTOM5 consortium (Andersson, Gebhard, et al. 2014). This yielded a single set of 30,439 autosomal enhancer coordinates. We assigned ages to enhancer sequences by intersecting their genomic coordinates with aged syntenic blocks using Bedtools v2.27.1 (Quinlan and Hall 2010). Syntenic blocks that overlapped at least 6 bp of an enhancer sequence (reflecting the minimum size of a TF binding site (Lambert et al. 2018) were considered when assigning the enhancer’s age and architecture. We considered enhancers with one age observed across its syntenic block(s) as “simple” enhancer architectures and enhancers overlapping syntenic blocks with different ages as “complex” enhancer architectures. We assigned complex enhancers ages according to the oldest block. Sequences without an assigned age were excluded from this analysis.

### cCRE enhancer data, age assignment, and architecture mapping

We considered HepG2 and K562 ENCODE3 candidate cis-regulatory elements (cCRE) enhancer loci annotated with proximal or distal enhancer-like signatures (pELS or dELS, with and without CTCF binding) (The ENCODE Project Consortium et al. 2020). This yielded 53,864 HepG2 and 46,188 K562 cCREs coordinates. As for eRNA, we assigned ages and architectures to enhancer sequences by intersecting their locations with hg38 syntenic blocks and evaluating the diversity of syntenic ages. Syntenic blocks that overlapped at least 6 bp of an enhancer sequence were considered when assigning the enhancer’s age and architecture. Complex enhancer architectures were defined as sequences with more than one age.

### MPRA activity data

MPRA activity data and tile coordinates as assayed by the SHARPR-MPRA approach (Ernst et al. 2016) were downloaded and filtered for “Enh”, “EnhF”, “EnhW”, and “EnhWF” ChromHMM annotations. All tiles were 295 base pairs in length. We intersected autosomal MPRA tile coordinates with syntenic blocks and assigned ages and architectures as described above for other enhancers.

### Genome-wide shuffles to determine expected background distributions

For FANTOM and cCRE enhancers, we performed 100 genomic shuffles of all regions in each dataset using Bedtools. These shuffled sets were matched to the chromosome and length distribution of the observed regions. Coding sequences and ENCODE blacklist regions were excluded (Amemiya, Kundaje, and Boyle 2019, https://www.encodeproject.org/annotations/ENCSR636HFF/). Each set of shuffled non-coding “background” genomic regions was then assigned ages and architectures with the same strategy used for the observed enhancers.

### TFBS density and enrichment

Coordinates for ENCODE3 ChIP-seq peaks for 119 and 249 transcription factors assayed in HepG2 and K562, respectively, were downloaded from the SCREEN ENCODE project (https://screen.encodeproject.org, last downloaded Feb. 14th, 2021). To assign TFBS to enhancer components, we intersected the 30 bp around the peak midpoint with simple and complex enhancer coordinates from the matching cell line. ChIPseq peaks overlapping enhancers by ≥6 bp were counted as overlapping and peak overlap counts were normalized by syntenic length to estimate the density of TFBS per base pair for each enhancer component.

For TFBS density and binding site enrichment, we only considered complex enhancers where TFBS overlapped enhancers. To correlate core and derived TFBS density, some complex enhancers have multiple derived sequences which complicates the comparison of core and derived TFBS density. Thus, for this analysis, we calculated TFBS density as the sum of TFBS sites divided by the sum of the length of derived or core regions. We observed similar result when considering pair-wise sytenic TFBS densities and summed core-derived TFBS densities (Fig S10, 11). For TFBS enrichment, we used regions matched on core and derived sequence ages to compare TFBS enrichment among sequences that emerged in the same evolutionary period. Per age TFBS enrichment in derived v. core regions was calculated as the number of TFBS peaks that bind these regulatory regions versus all other TFBS loci that bind regulatory regions in that evolutionary period. Fisher’s exact test was used to compute the odds ratios and P-values, which were corrected for multiple hypothesis testing to control the false discovery rate at 5% using the BenjaminiHochberg procedure

### 1000 genomes variant density and minor allele frequency analyses

Genetic variants from 2504 diverse humans were downloaded from the 1000G project phase3 (shapeit2 mvncall integrated v5a release 20130502). We intersected all variants with FANTOM enhancers and stratified by core and derived regions. Variant density was estimated as the number of SNPs overlapping a syntenic block divided by the length of the syntenic block. Singletons, i.e. alleles observed only once in a single individual, were removed from this analysis.

### LINSIGHT purifying selection estimates

Pre-computed LINSIGHT scores were downloaded from http://compgen.cshl.edu/~yihuang/LINSIGHT/. LINSIGHT provides per base pair estimates of the probability of negative selection (Huang, Gulko, and Siepel 2017). We intersected FANTOM enhancers with LINSIGHT bp scores to determine the levels of constraint on bases within core and derived sequences.

### eQTL enrichment

The enrichment for GTEx eQTL from 46 tissues (last downloaded July 23rd, 2019) in core and derived enhancer sequences was tested against a null distribution determined by permutations. Enhancer sequences were shuffled to create 500 sets of random genomic regions matched on lengths and chromosome. Median fold-change was calculated as the number of eQTLs overlapping enhancer sequence components (i.e., core or derived) compared with the appropriate random sets. Confidence intervals (CI = 95%) were generated by 10,000 bootstraps. P-values were corrected for multiple hypothesis testing by controlling the false discovery rate (FDR) at 5% using the Benjamini-Hochberg procedure.

## Data availability

### Sequence age datasets

- Hg19 syntenic age data (including aged FANTOM eRNAs) underlying this article are available in Zenodo, at https://dx.doi.org/10.5281/zenodo.4618495
- Hg38 syntenic age data underlying this article are available in Zenodo, at https://doi.org/10.5281/zenodo.5809634
- HepG2 and K562 aged cCRE sequences underlying this article are available in Zenodo, at https://doi.org/10.5281/zenodo.5809629

### Datasets derived from sources in the public domain

- FANTOM5 eRNAs (Andersson, Gebhard, et al. 2014) - http://slidebase.binf.ku.dk/human_enhancers/
- ENCODE cCREs and TFBS ChIP-seq (The ENCODE Project Consortium et al. 2020) - https://screen.encodeproject.org
- HepG2 and K562 MPRAs (Ernst et al. 2016) - GSE71279
- Hg19 46-way vertebrate species multiz alignment - https://hgdownload.soe.ucsc.edu/gbdb/hg19/multiz46way/
- Hg38 100-way vertebrate species multiz alignment - https://hgdownload.soe.ucsc.edu/gbdb/hg38/multiz100way/
- LINSIGHT (Huang, Gulko, and Siepel 2017) - http://compgen.cshl.edu/LINSIGHT/LINSIGHT.bw

### Source code is openly available at

https://github.com/slifong08/enh_ages

