## Supplemental Material for "Function and constraint in enhancers with multiple evolutionary origins"

### Supplemental: Evolutionarily younger segments of enhancers have distinct functional properties

December 24, 2021

14 Age-matched derived regions are under lower purifying selection pressures across core ages. 15
15 Derived regions have higher minor allele frequencies than core regions across human populations. . . . . . 16
16 Derived regions have higher SNP densities than adjacent core regions . . . . . 17
17 ChIP-seq TFBS binding frequency in core and derived regions of HepG2 and K562 complex enhancers from ENCODE . . . . . 18
18 GC density in FANTOM enhancer and promoter regions . . . . . 18

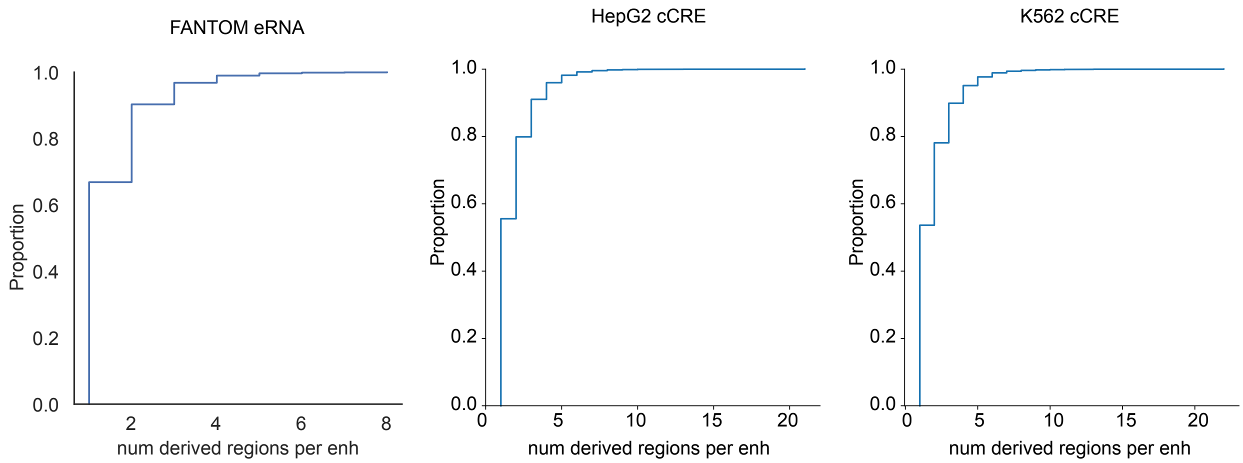

**Figure 1: Number of derived regions per complex enhancer**

**Most complex enhancers have one derived region.** Cumulative distribution plots show the number of derived regions as proportion of the total complex enhancer sequences for FANTOM5 eRNA (left, N = 10851), HepG2 cCREs from ENCODE (middle, N = 27289) and K562 cCREs from ENCODE (right, N = 24415). Complex enhancers have a median of one derived region across datasets.

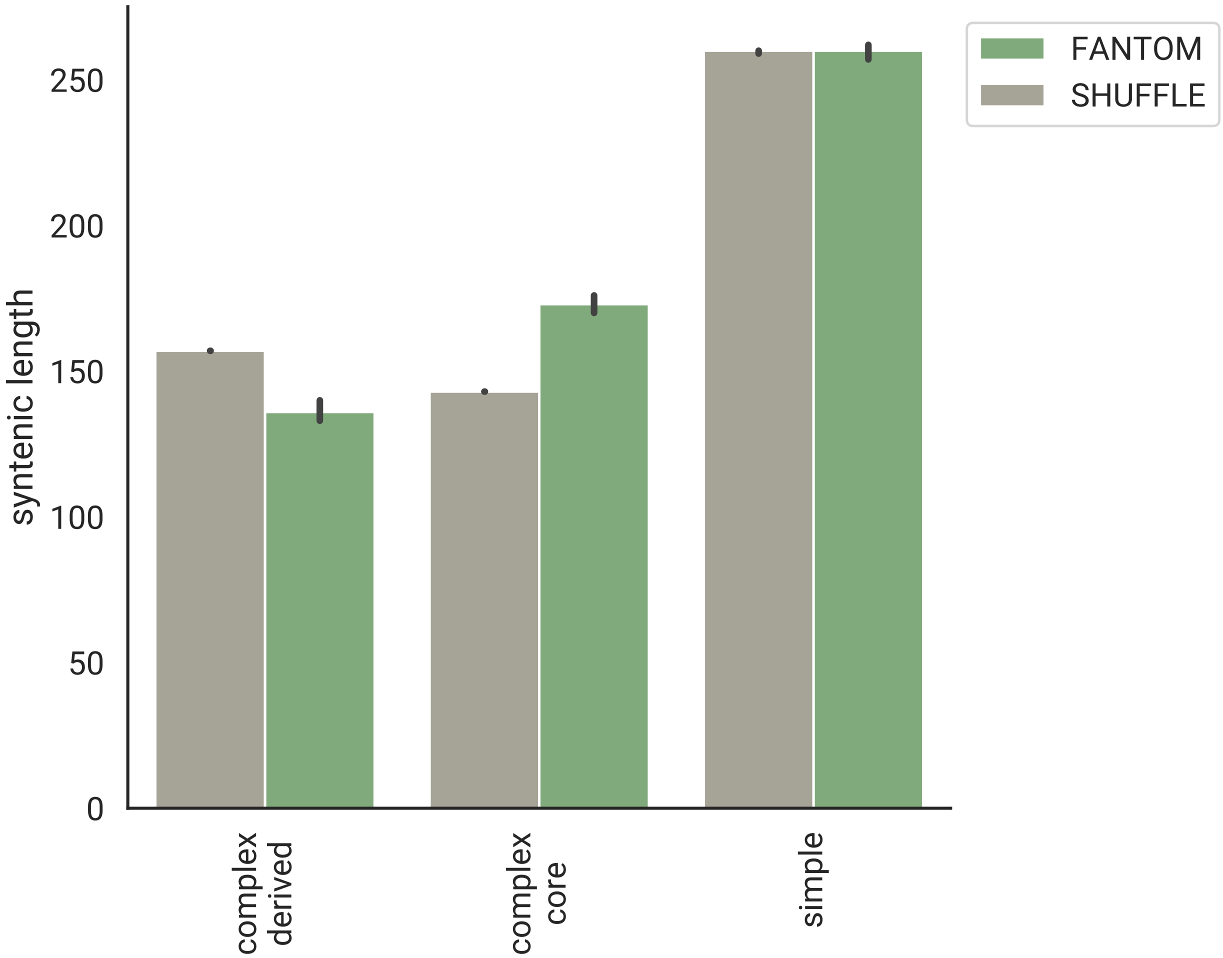

**Figure 2: Sequence lengths of derived, core, and simple transcribed enhancers**

**Derived regions are also shorter than expected** from 100 sets of length-, chromosome-, and architecture-matched random non-coding regions (left; median 136 bp derived v. 157 bp shuffled, p = 1.4e-46). Core sequences in complex enhancers are longer than 100x non-coding, chromosome-matched shuffled background cores (right; median 173 bp core v. 143 bp shuffle core, p = 2.4e-75). Sample size is annotated for each bar

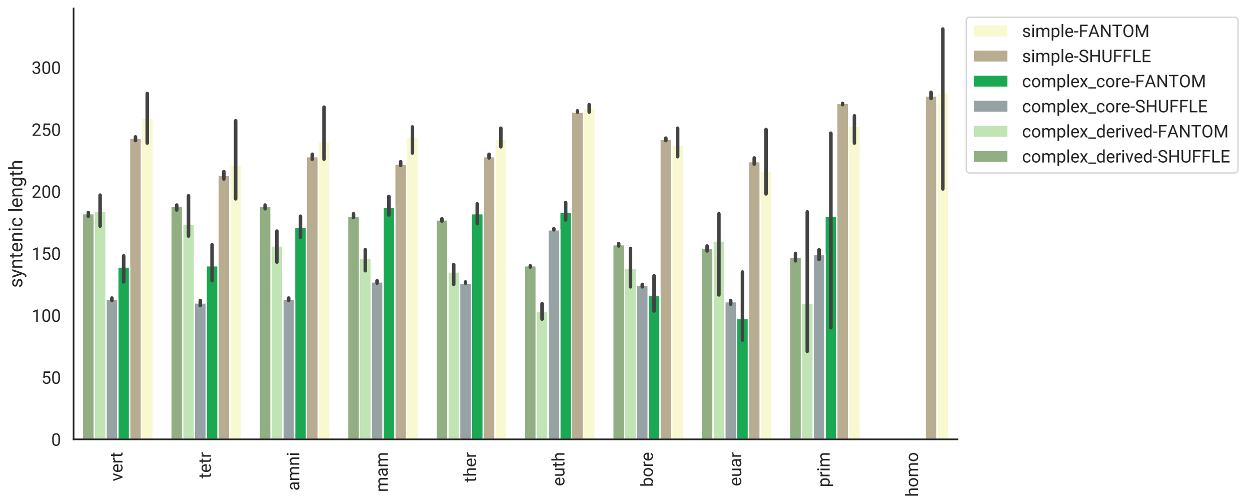

**Figure 3: Lengths of derived, core, and simple enhancers versus expectation, stratified by core age**

Derived, core and simple sequence lengths stratified by core age (x-axis) and compared with 100x shuffled sequences matched on core sequence age and architecture. Derived sequences are shorter than expected at every age except those with Vertebrate cores. Core sequences from the Eutherian ancestor and older are longer than expected.

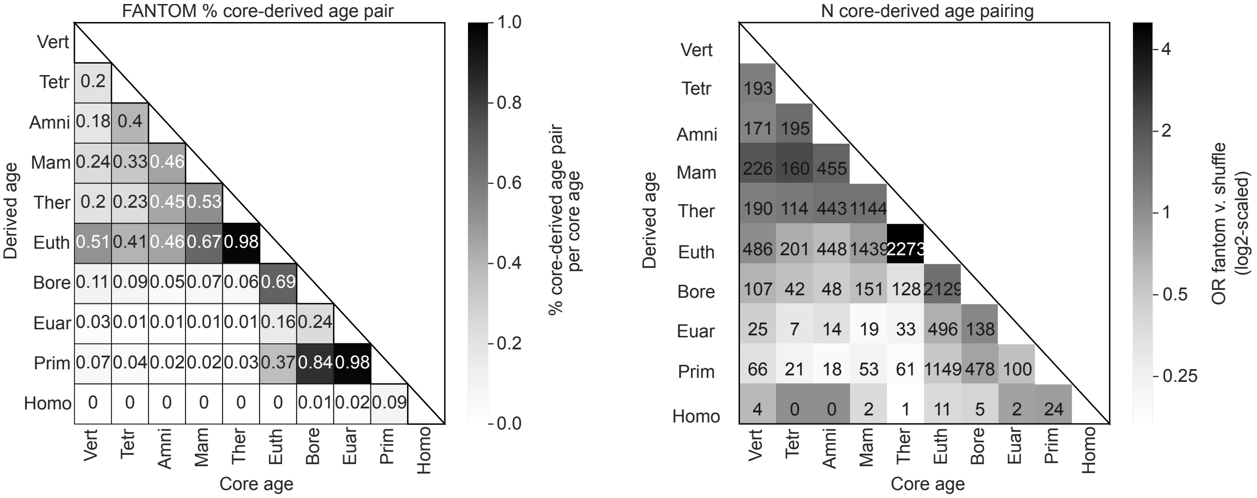

**Figure 4: Frequency and count of core-derived sequence age pairs in FANTOM**

Frequency (left) and count (right) of core-derived age pairs across complex FANTOM enhancers. Shading in the frequency plot (left) reflects the percentage of age-pairs within a single core age. Cores may have more than one derived sequence of a different age, thus the sum of the columns can be greater than one. Shading in the count plot (right) reflects the enrichment of the core-derived age pair compared with shuffled expectation shown in Figure 3.

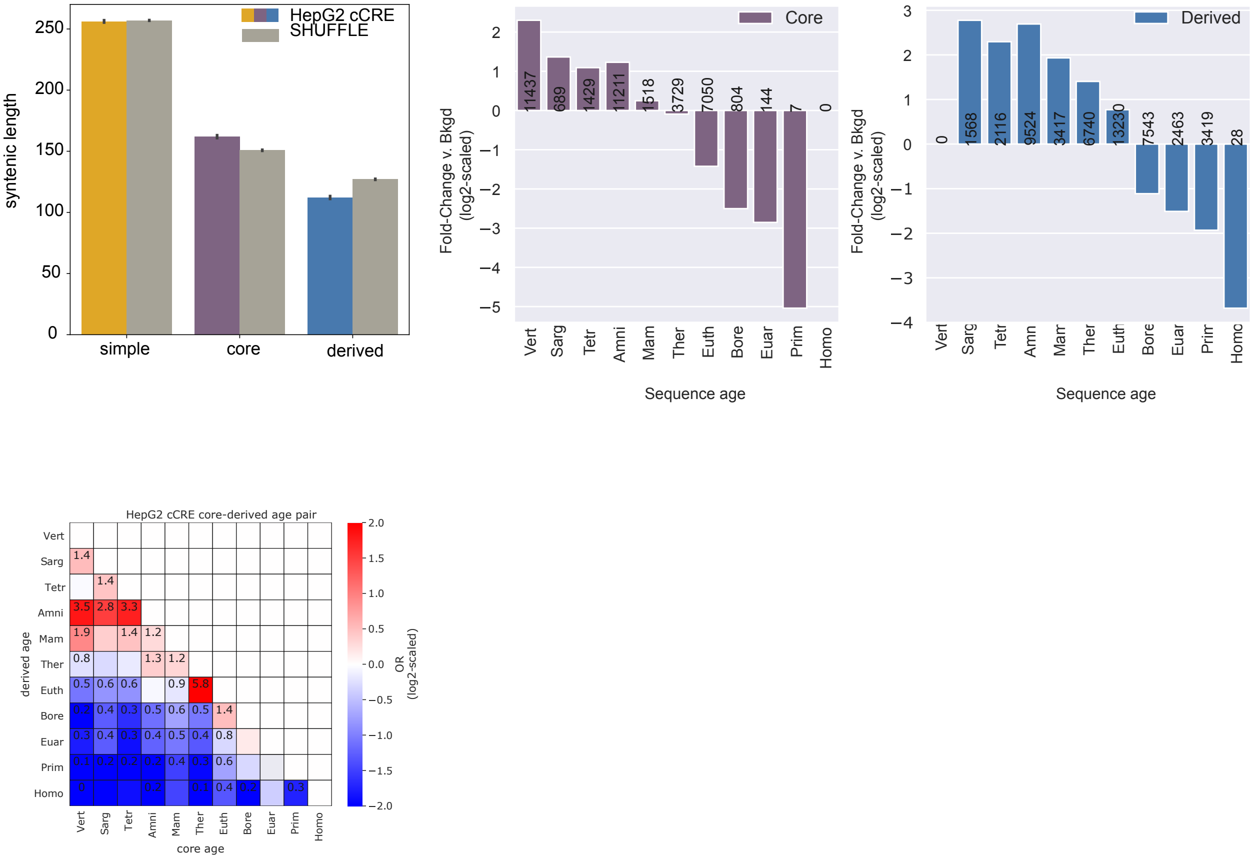

**Figure 5: Core and derived evolutionary features in complex HepG2 cCREs recapitulate evolutionary features in FANTOM eRNAs**

Derived regions constitute a sizeable portion of complex HepG2 cCREs (N = 27,789 cCREs), are shorter (top left) and older (top right) than expected compared to shuffled complex enhancer architectures (N = 1,047,557). Core sequences from the Mammalian ancestor and older are enriched for derived sequences from the Therian ancestor and older compared with shuffled expectation of core-derived age pairs. These core sequences are also depleted of sequences younger than the Therian ancestor. Core sequences are enriched for the nearest, younger phylogenetic neighbor. Odds ratio of significantly enriched age-pairs (FDR$<$ 0.05) are annotated.

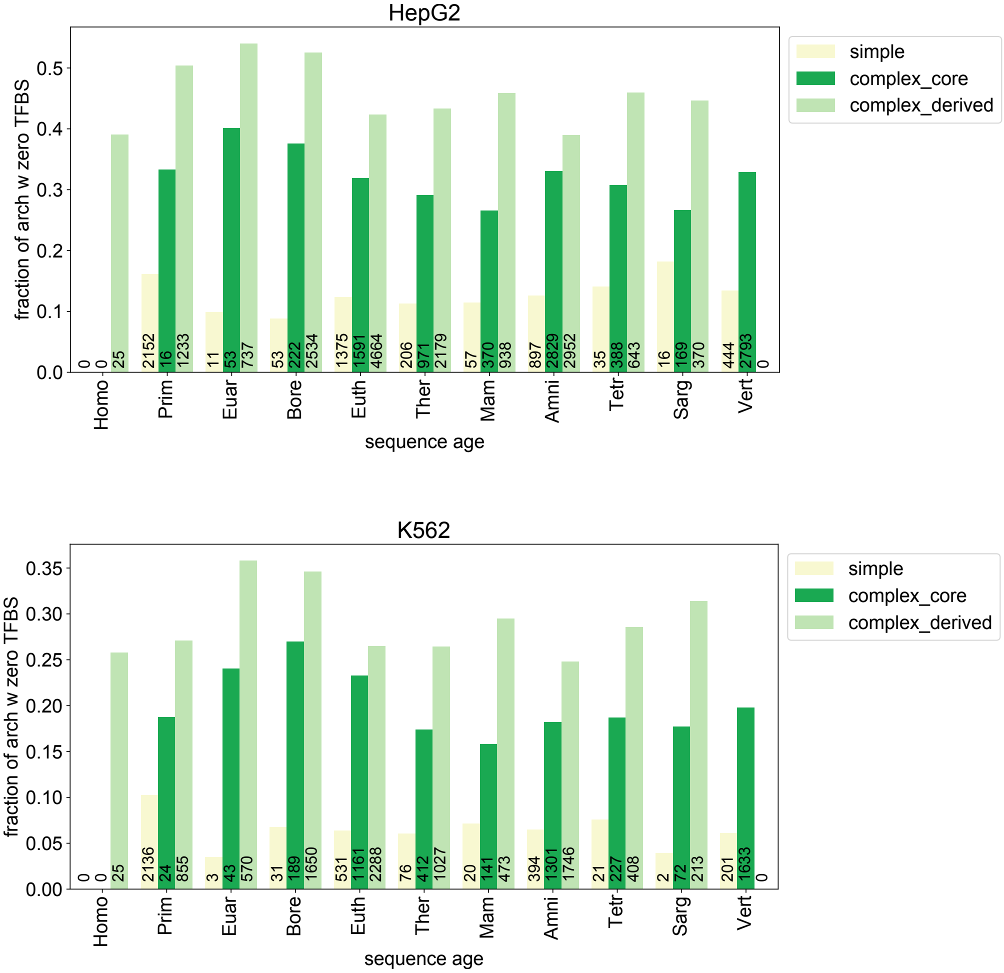

**Figure 6: Regions with no TFBS ChIP-seq binding are observed across ages**

Similar proportions of derived, core, and simple enhancer sequences have no evidence of TFBS binding within sequence ages in HepG2 and K562 cCREs. K562 cell models generally have fewer elements that do not overlap TFBS, likely because more TFBS ChIP-seq assays have been performed in K562 cells compared with HepG2 cells (249 v. 119 assays, respectively). Enhancer regions are binned according to their syntenic sequence ages. Frequency is calculated as the percent of regions that do not overlap TFBS ChIP-seq peaks within each sequence age and region category. HepG2 is shown above and K562 is shown below. Number of regions with zero TFBS overlap is annotated for each bar.

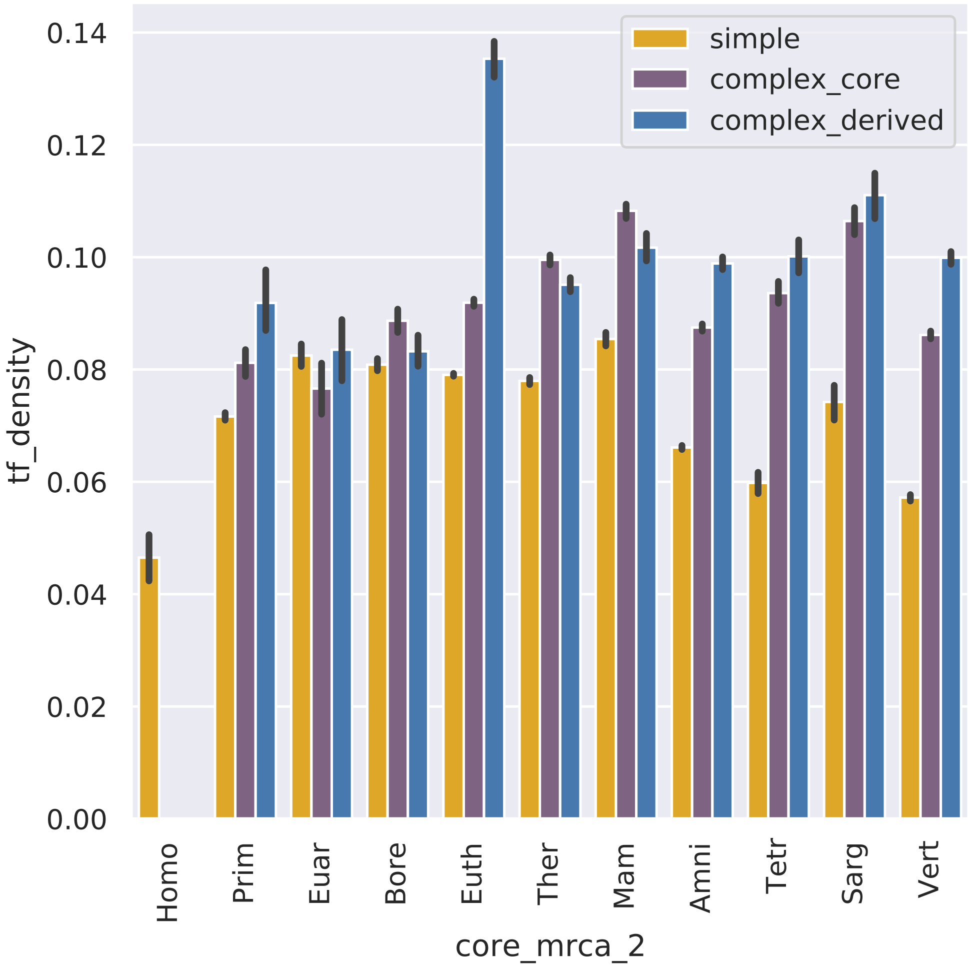

**Figure 7: Transcription factor binding site density is similar across ages in HepG2 cCREs**

Simple, core, and derived sequences are stratified by core age on the x-axis. TFBS density per architecture and age was measured and plotted on the y-axis.

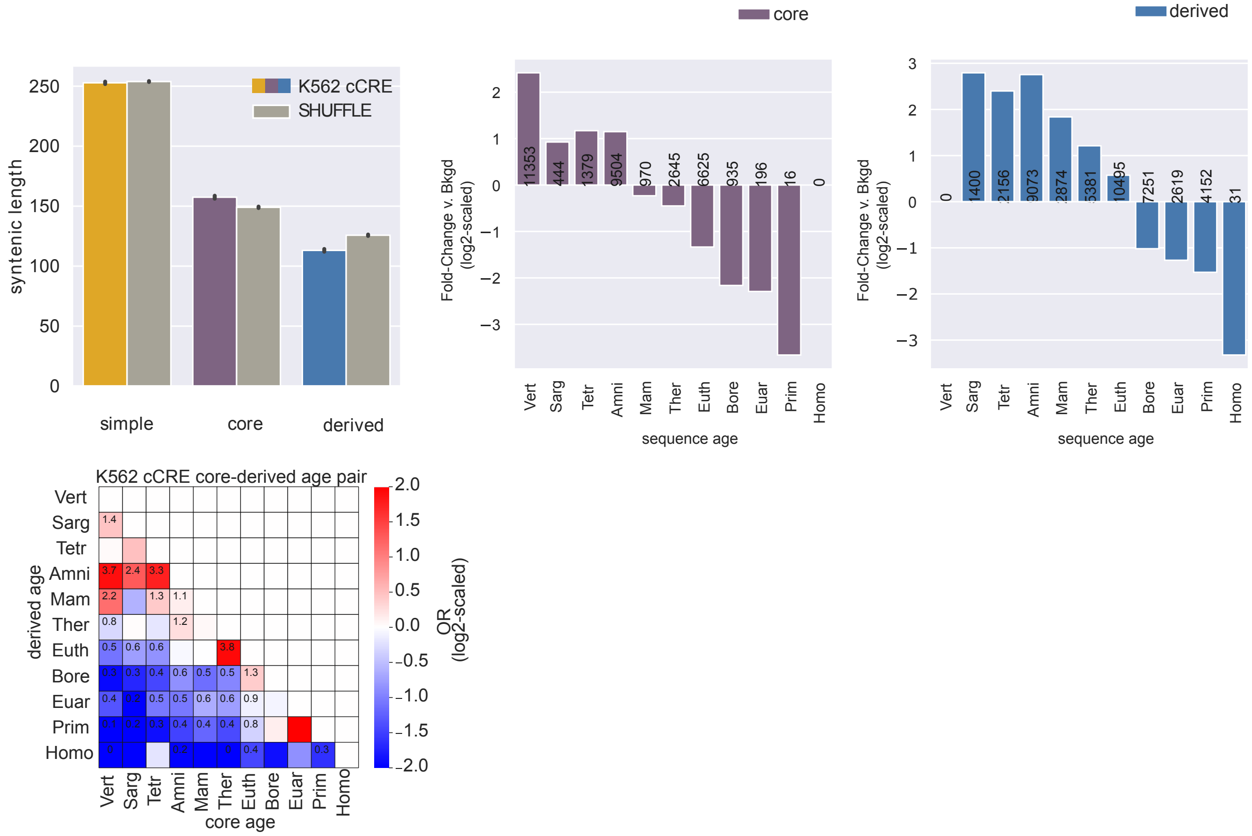

**Figure 8: Core and derived evolutionary features in complex K562 cCREs recapitulate evolutionary features in FANTOM eRNAs**

Derived regions constitute a sizeable portion of complex K562 cCREs (N = 24,415 cCREs), are shorter (top left) and older (top right) than expected compared to shuffled complex enhancer architectures (N = 473,387 cCREs). Core sequences from the Amniota ancestor and older are enriched for derived sequences from the Mammalian ancestor and older compared with shuffled expectation of core-derived age pairs. These core sequences are also depleted of sequences younger than the Mammalian ancestor. Core sequences are enriched for the nearest, younger phylogenetic neighbor. Odds ratio of significantly enriched age-pairs (FDR$<$ 0.05) are annotated.

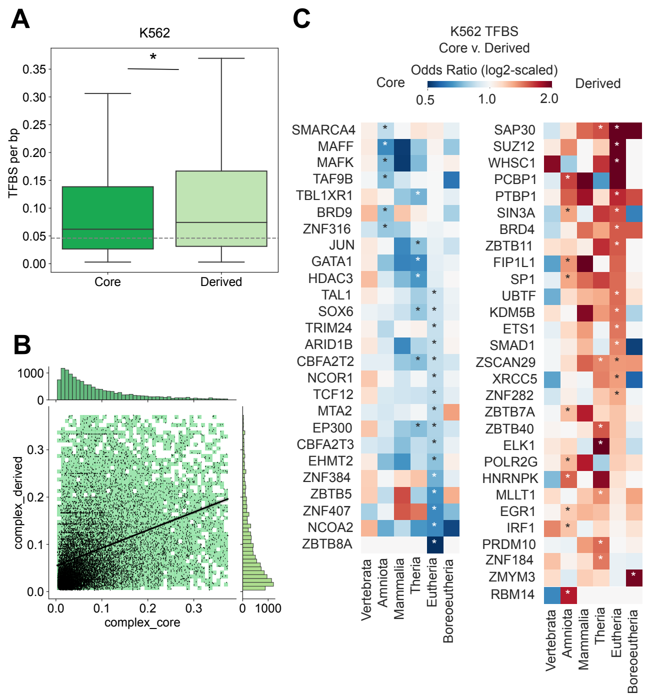

**Figure 9: Derived regions have high transcription factor binding site densities and bind different transcription factors compared to core regions in K562 cells**

**(A)** Derived regions (N = 23868) have higher TFBS densities than core regions (N = 20997) (0.074 derived v. 0.062 core TFBS per base pair, Mann Whitney-U p = 3.5e-52). Simple enhancer TFBS density is lower than core and derived regions (0.05 TFBS per base pair) **(B)** TFBS density is positively correlated between core-derived sequence pairs within complex enhancers with evidence of TF binding in both core and derived regions (N = 14142). Color intensity represents the density of core-derived pairs, and the black line is a linear regression fit (slope=0.39, intercept=0.056, r=0.39, p $<$2.2e-238, stderr=0.008; outliers ($>$95th percentile) are not plotted for ease of visualization. **(C)** Derived and core regions of the same age are enriched for binding of different TFs and enrichment patterns are generally consistent across ages. TFBS enrichment for each age was tested using Fisher's exact test; only TFs with at least one significant enrichment (FDR $<$ 0.1) are shown. Vertebrate, Sarcopterygii, and Tetrapod enhancer ancestors were grouped into "Vert+". Boreotherian, Euarchontoglires, and Primate enhancer ancestors were grouped into "Bore+".

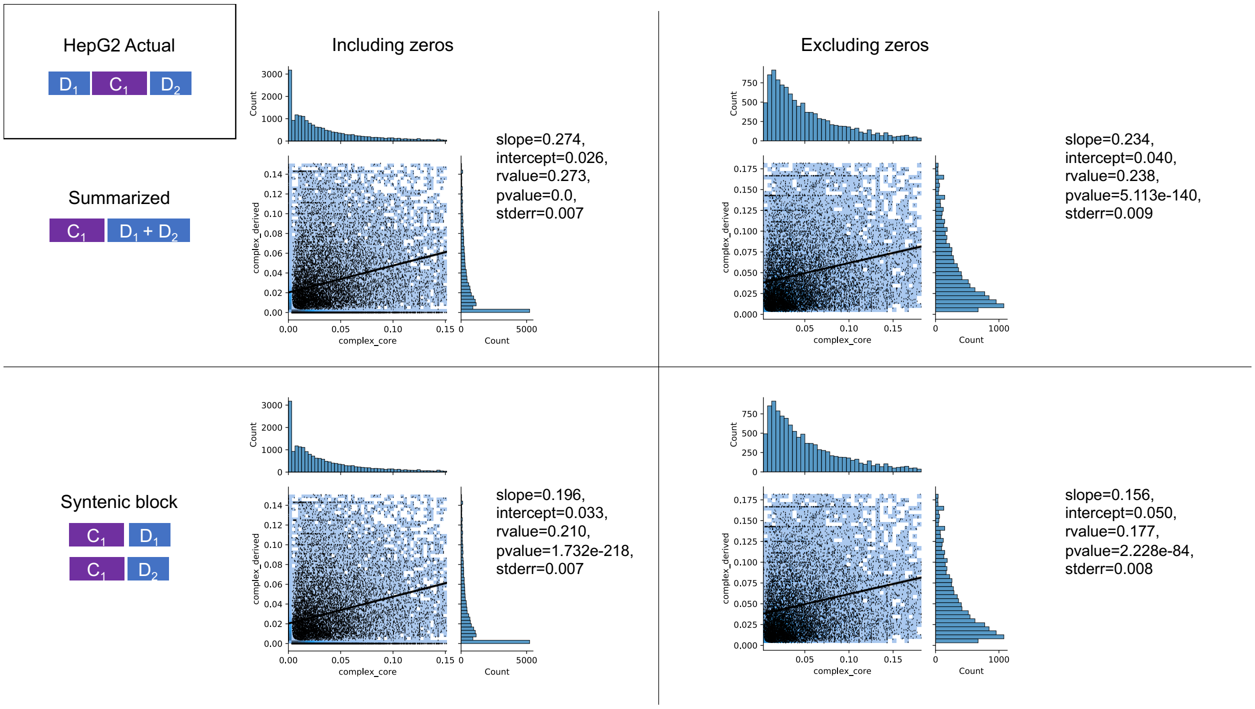

**Figure 10: High TFBS density in core regions correlates with high TFBS density in derived regions within the same HepG2 enhancer sequence**

We evaluated TFBS density correlations between core and derived sequences of the same enhancer in HepG2 cCREs. TFBS density of core and matched-derived regions per enhancer are plotted on the X- and Y-axis, respectively. Actual enhancers can have more than one core or derived region, so we evaluated our data using two different approaches. In the first (upper) we summarized TFBS density across multiple core and derived regions by summing TFBS density and syntenic length into core and derived groups and quantifying TFBS density in summarized core and derived regions per enhancer sequence. In the second (lower), we quantified TFBS density for every core and derived syntenic region and compared all possible pairs of core and derived syntenic TFBS densities per enhancer. We applied two different thresholds for evaluating TFBS density in core versus matched derived regions; one threshold allowed for regions with no evidence of TFBS in core or derived sequence, but not both (left, “zeros included”), while the other threshold required that TFBS binding was detected in both core and derived sequences within an enhancer (right, “zeros excluded”). Linear regression models were fit for each dataset and model features are annotated for each analysis. Histograms (right and above) display distributions of core and derived TFBS density per analysis.

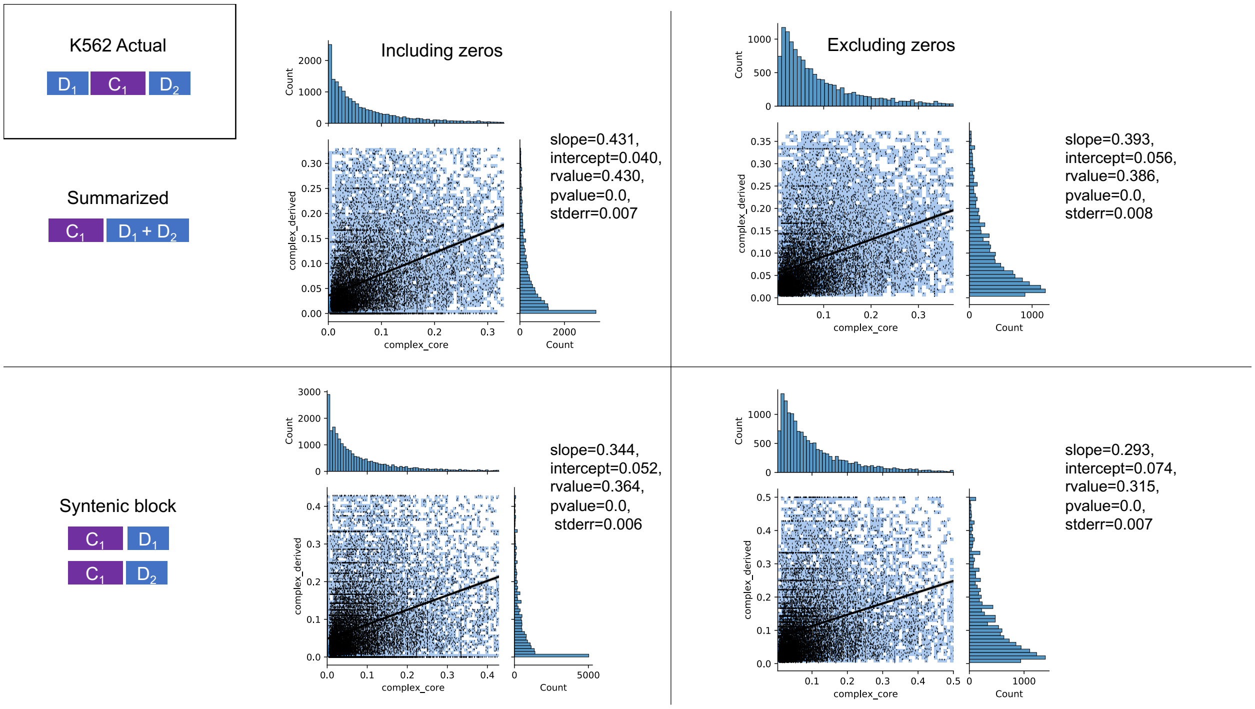

**Figure 11: High TFBS density in core regions correlates with high TFBS density in derived regions within the same K562 enhancer sequence**

We evaluated TFBS density correlations between core and derived sequences of the same enhancer in K562 cCREs. TFBS density of core and matched-derived regions per enhancer are plotted on the X- and Y-axis, respectively. Actual enhancers can have more than one core or derived region, so we evaluated our data using two different approaches. In the first (upper) we summarized TFBS density across multiple core and derived regions by summing TFBS density and syntenic length into core and derived groups and quantifying TFBS density in summarized core and derived regions per enhancer sequence. In the second (lower), we quantified TFBS density for every core and derived syntenic region and compared all possible pairs of core and derived syntenic TFBS densities per enhancer. We applied two different thresholds for evaluating TFBS density in core versus matched derived regions; one threshold allowed for regions with no evidence of TFBS in core or derived sequence, but not both (left, “zeros included”), while the other threshold required that TFBS binding was detected in both core and derived sequences within an enhancer (right, “zeros excluded”). Linear regression models were fit for each dataset and model features are annotated for each analysis. Histograms (right and above) display distributions of core and derived TFBS density per analysis.

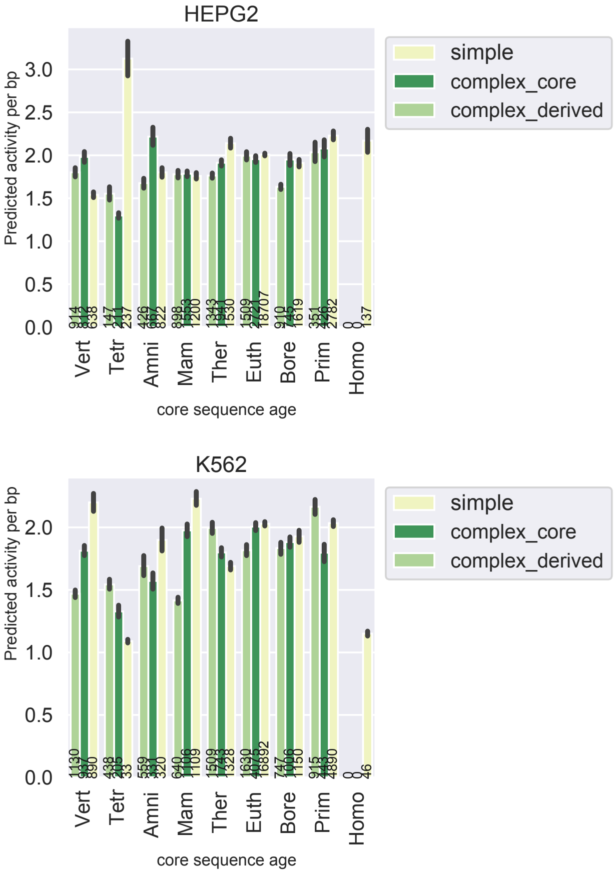

**Figure 12: MPRA activity is similar across sequence ages and simple, core, or derived contexts**

MPRA predicted activity per bp from Ernst 2016 is similar across ages in K562 and HepG2 cells. Here, predicted activity per bp scores are stratified by core sequence age and simple, core, or derived category. Cell line models and N bp are annotated per bar.

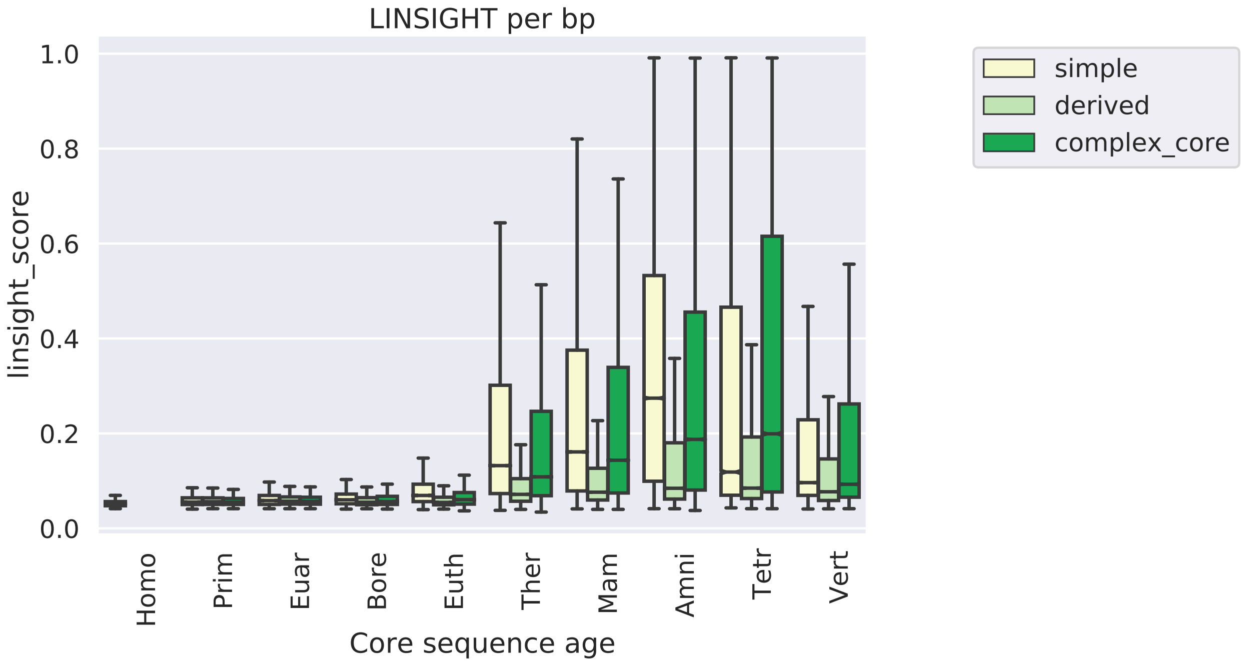

**Figure 13: Age-matched derived regions are under lower purifying selection pressures across sequence ages.**

Derived regions are under lower purifying selection pressures across sequence ages than core or simple FANTOM enhancers. Per basepair LINSIGHT purifying selection scores were estimated in Huang et at 2017. Stratified by core sequence age, derived sequences have lower LINSIGHT scores than adjacent core sequences for all core sequences older than Boreotherian. Number of measurements is annotated per bar.

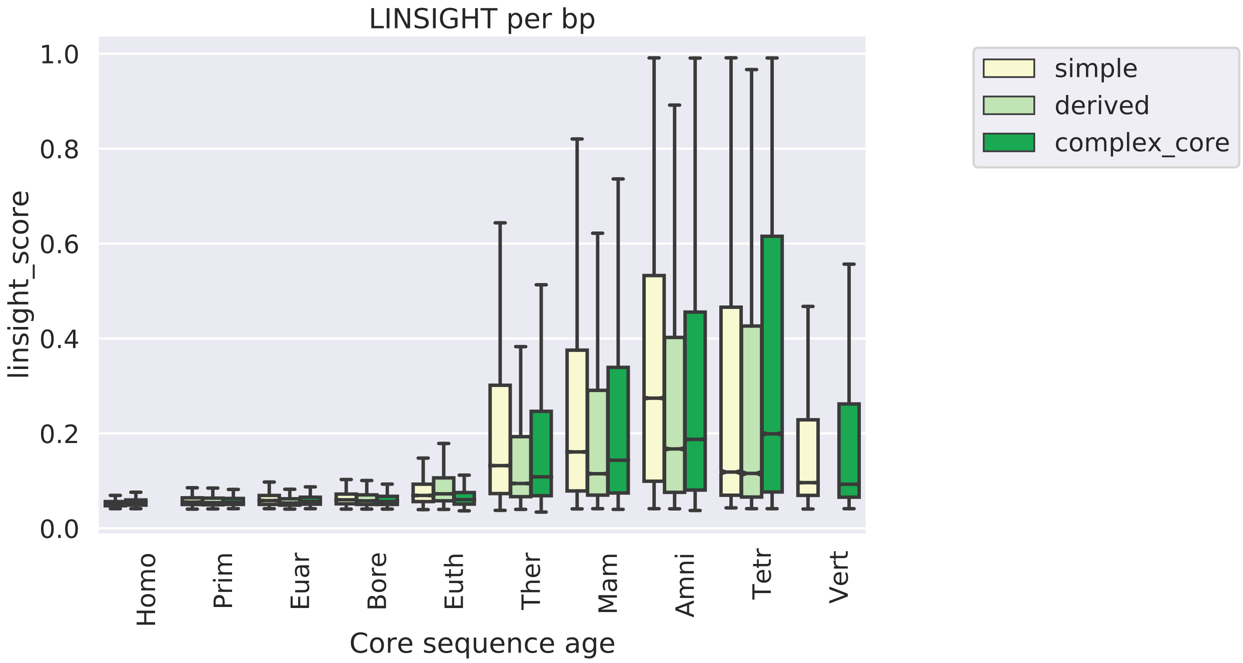

**Figure 14: Age-matched derived regions are under lower purifying selection pressures across core ages.**

Per basepair LINSIGHT purifying selection scores were estimated in Huang et at 2017 with FANTOM eRNA. Stratified by syntenic sequence age, derived sequences have lower LINSIGHT scores than age-matched core sequences for all sequences older than Eutherian. Number of measurements is annotated per bar.

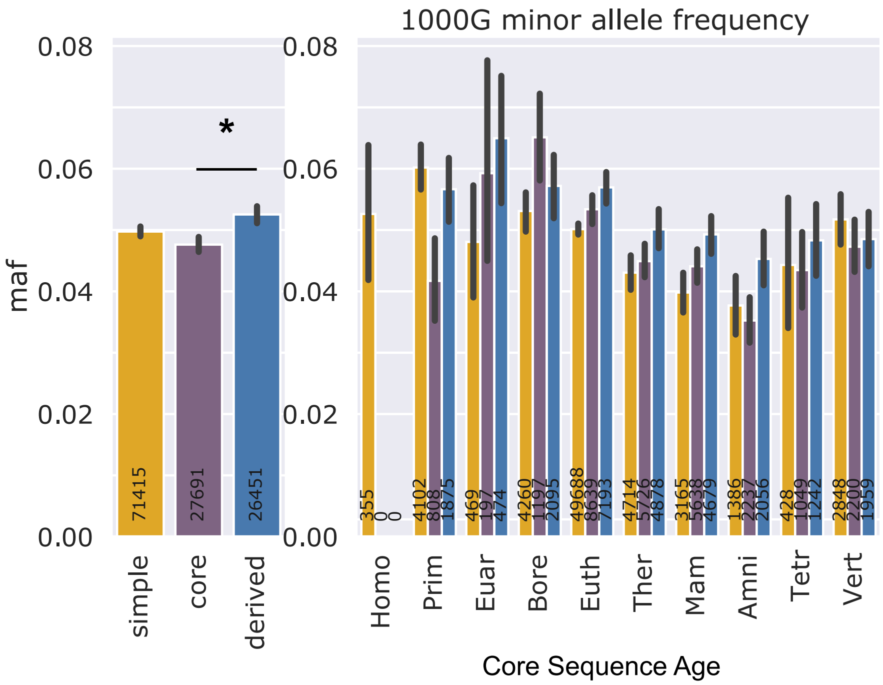

**Figure 15: Derived regions have higher minor allele frequencies than core regions across human populations.**

Global minor allele frequencies from 1000 Genomes were intersected with FANTOM enhancer components. Singletons were removed. Derived region minor allele frequencies are slightly higher than core region minor allele frequencies (right, mean 0.053 derived v. 0.048 core, derived v. core p = 4.9e-12). Minor allele frequencies stratified by core age and architecture show that derived regions have consistently higher minor allele frequencies compared to core regions at every ancestral origin except Boreotherian. Number of SNPs is annotated per bar.

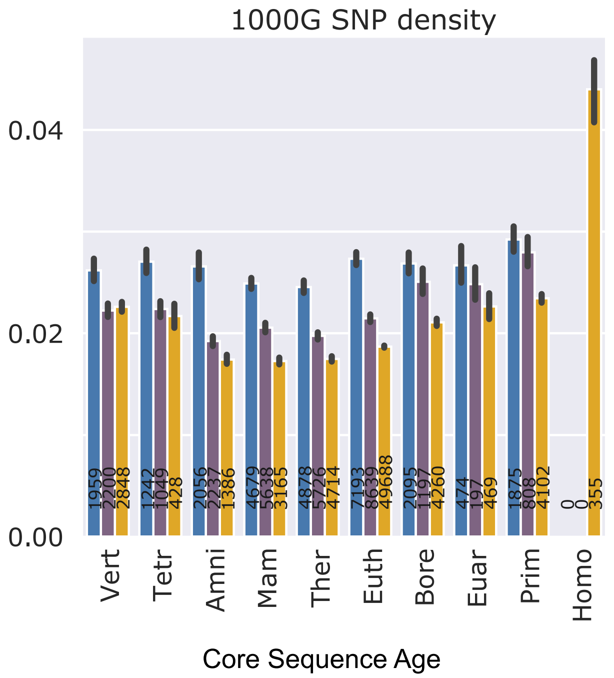

**Figure 16: Derived regions have higher SNP densities than adjacent core regions**

SNP densities from 1000G were calculated as the number of SNPs in a region divided by the syntenic length. Densities were then stratified by architecture and core age. Number of SNP is annotated per bar.

| Cell line | arch | Count zero TF overlap | Total counts | freq zero TF overlap | freq TF overlap |
|---|---|---|---|---|---|
| HepG2 | derived | 23955 | 44222 | 0.54 | 0.46 |
| HepG2 | core | 9859 | 29832 | 0.33 | 0.67 |
| HepG2 | simple | 3390 | 26624 | 0.13 | 0.87 |
| K562 | derived | 16572 | 40110 | 0.41 | 0.59 |
| K562 | core | 5731 | 26728 | 0.21 | 0.79 |
| K562 | simple | 1587 | 22768 | 0.07 | 0.93 |

**Table S**

**Figure 17: ChIP-seq TFBS binding frequency in core and derived regions of HepG2 and K562 complex enhancers from ENCODE**

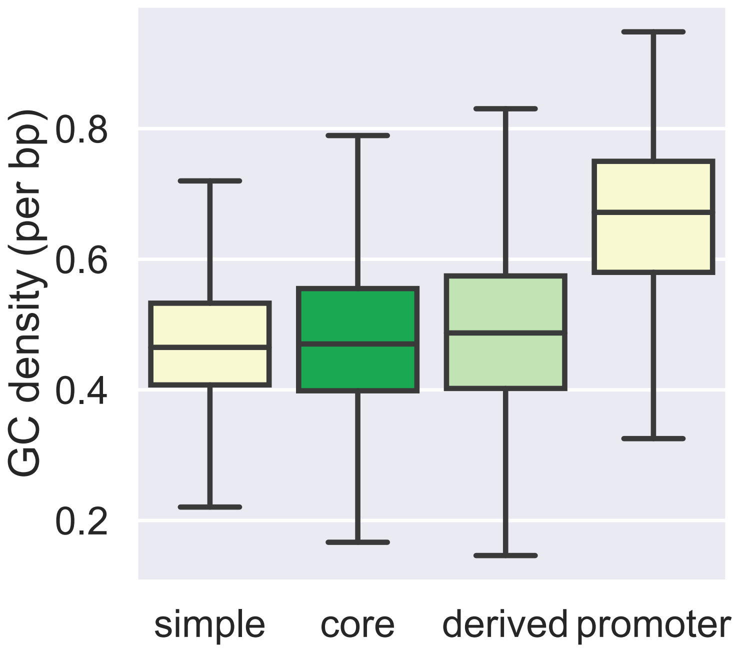

**Figure 18: GC density in FANTOM enhancer and promoter regions**

GC density was calculated across FANTOM enhancers and promoters as the number of G or C bases divided by the length. Non-exonic enhancers have lower GC density than promoters (N =13781). Derived regions (N = 15357) have slightly higher GC density than core regions (N = 11489) (median 0.49 derived v. 0.47 core GC density; MWU p = 1.7e-12). Simple enhancers (N= 20087) have similar GC density to enhancer cores (median 0.47 GC density)
